# Activation and inactivation pathways of a p53-like transcription factor govern lipid homeostasis in yeast

**DOI:** 10.64898/2026.08.31.748402

**Authors:** Neng Wan, Jeremi Kuklewicz, Joao A. Paulo, Christopher Nardone, Steven P. Gygi, Tom A. Rapoport

## Abstract

Membrane fluidity depends on unsaturated acyl chains that are generated in Saccharomyces cerevisiae by the desaturase Ole1, whose expression is primarily under the control of the transcription factor Mga2. Here, we show that the endoplasmic reticulum-anchored Mga2 precursor is ubiquitinated by the E3 ligase Rsp5 and then processively degraded by the proteasome until proteolysis stalls at a defined site, releasing a soluble fragment that enters the nucleus and activates Ole1 transcription. Unexpectedly, Mga2 contains a DNA-binding domain and a trans-activation-like segment structurally and functionally related to those of the human tumor suppressor p53. The mature transcription factor is degraded in the nucleus in a DNA binding-dependent manner; blocking this degradation causes unsaturated acyl chains to accumulate in lipid droplets, a detoxification response required for cell viability. These findings define the pathways that activate and inactivate Mga2, and uncover an evolutionary connection between the yeast lipid homeostasis regulator Mga2 and p53.

## INTRODUCTION

All cells must maintain appropriate membrane fluidity by controlling the degree of unsaturation of acyl chains in their lipids. Too few unsaturated acyl chains compromise phospholipid bilayer flexibility, while an excess causes membrane leakiness. The enzymes that generate unsaturated fatty acids are therefore tightly regulated. In mammals, double bonds are introduced by the fatty acid desaturases SCD1, whose transcription is regulated by the transcription factor SREBP1 (*1*). In Saccharomyces cerevisiae, Ole1, a homolog of SCD1, is the sole enzyme that generates unsaturated acyl chains. Ole1 is a multi-spanning Δ9-desaturase in the endoplasmic reticulum (ER) membrane, which uses molecular oxygen as a co-substrate (*2*). Ole1 transcription is repressed at high levels of unsaturated fatty acids (*3, 4*) and induced under low-oxygen conditions to compensate for the reduced enzymatic activity of Ole1 in hypoxia (*4, 5*). This regulation is mediated by two related transcription factors, Mga2 and Spt23 (*6–10*). Yeast cells tolerate the loss of either Mga2 or Spt23 individually, but deletion of both is lethal unless exogenous oleate is supplied (*7, 8*), consistent with the essential role of Ole1 in generating unsaturated fatty acids.

Like SREBP1, Mga2 and Spt23 are synthesized as membrane-bound precursors that are proteolytically processed into soluble mature transcription factors. However, the actual SREBP1 and Mga2 pathways are quite different. The precursor of SREBP1 is moved to the Golgi and processed by two distinct proteases in a pathway that has been extensively studied (*11*). In contrast, the precursors of Mga2 and Spt23 are proteolytically processed in the endoplasmic reticulum (ER) by a pathway that is poorly understood. Both proteins are synthesized as single-pass membrane protein precursors of ∼120 kDa (p120). Processing of p120 into a soluble cytosolic fragment of ∼90 kDa (p90) allows the mature transcription factor to translocate to the nucleus and activate Ole1 transcription (*10*). Unsaturated fatty acids are thought to suppress this processing by inducing rotation of the TM segments within the p120 dimer (*12*). The mechanism of p120-to-p90 conversion is unclear. The ubiquitin ligase Rsp5 has been implicated in this process (*8*), but its precise role is contested, as some studies found Rsp5 to be dispensable for Mga2 precursor processing (*13, 14*). Cleavage of p120 to p90 requires the 26S proteasome (*15*), yet why the proteasome degrades p120 only partially is unclear. One model proposes that the proteasome initiates degradation from ubiquitination sites near the TM segment and proceeds toward the N-terminus until stalled by a tightly folded domain (*16*). Analogous partial degradation occurs during NF-κB processing, where proteasome stalling depends on a glycine-rich region that impairs grip by the ATPase subunits (*17, 18*); however, no equivalent glycine-rich sequence is present in Mga2 or Spt23, and the precise stalling site has not been identified.

The mature p90 fragment activates Ole1 transcription through two promoter elements: a fatty acid-regulated region (FAR)(*3*) and a low-oxygen response element (O_2_R/LORE)(*19–21*). How p90 mechanistically drives transcription remains unknown, because Mga2 and Spt23 were thought to contain no classical DNA binding domains (*10*). p90 has also been reported to be unstable (*8*), but whether this degradation occurs through the proteasome remains unresolved, as prior studies are confounded by the proteasome dependence of p120 processing (*22*). The biological relevance of p90 turnover has similarly not been demonstrated.

Here we resolve several of these outstanding questions. We show that the Rsp5 ubiquitin ligase modifies p120, which is then processively degraded by the proteasome to generate p90; proteasomal movement is arrested at a defined distance from the IPT domain. Strikingly, the resulting p90 fragment harbors a DNA-binding domain and a trans-activation-like segment that are structurally and functionally homologous to those of the mammalian tumor suppressor p53 — the “guardian of the genome” — a relationship that had not previously been recognized. Like p53, p90 is itself subject to proteasomal degradation, which we show occurs in the nucleus and depends on DNA binding. Failure to degrade p90 leads to toxic accumulation of unsaturated acyl chains, which are sequestered into lipid droplets as a detoxification mechanism essential for cell viability. Together, these findings define the pathways that activate and inactivate Mga2 and reveal an unexpected evolutionary relationship between a yeast transcription factor governing lipid homeostasis and the mammalian tumor suppressor p53.

## RESULTS

### Processing of p120 to p90

Although Mga2 and Spt23 are functionally redundant, Mga2 is reported to be present at higher levels than Spt23 (*23*). We confirmed this conclusion by tagging both proteins at their N-terminus with a hemagglutinin (HA) epitope and performing immunoblotting with HA antibodies (fig. S1A). Both the p120 precursor and the p90 fragment of Mga2 were more abundant than the corresponding species of Spt23. Mga2 was also more important for the expression of Ole1 mRNA and Ole1 protein, as shown by analyzing yeast strains that lack Mga2 or Spt23 (fig. S1, B and C). We therefore focused our analysis on Mga2.

The Mga2 precursor contains several domains predicted by AlphaFold (AF-P40578-F1): an unstructured N-terminal segment, a β-strand-rich domain (we now refer to as DNA-binding domain (DBD), see below), a dimerization domain (IPT), an ankyrin-repeat domain (Ank), a transmembrane segment (TM), and a C-terminal luminal domain (**Fig. 1A**). The IPT and Ank domains are similar to those in the mammalian transcription factor NF-κB, and as in NF-κB (*24, 25*), the IPT domain is thought to mediate precursor dimerization (*9*).

**Fig. 1.**
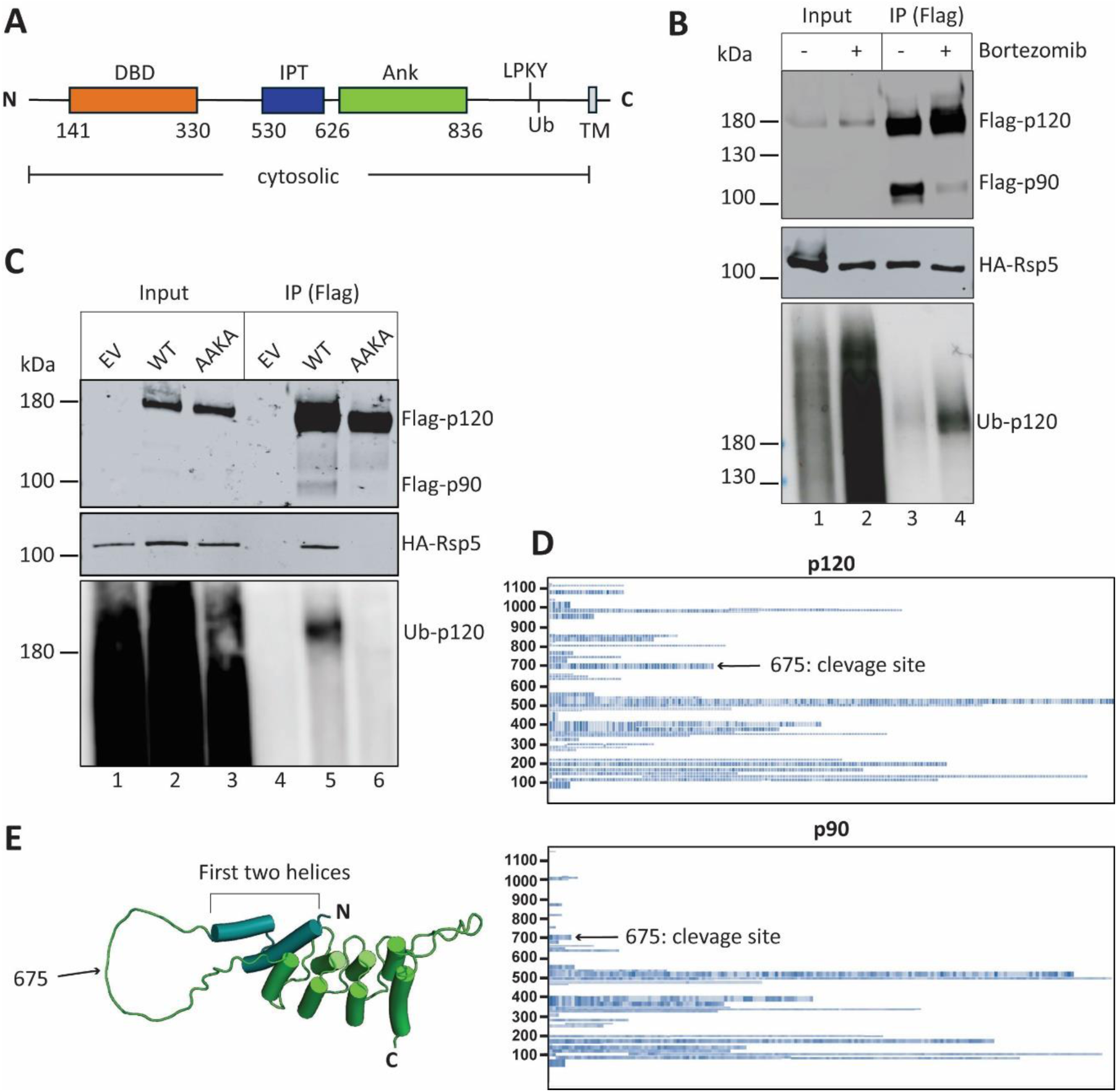
Processing of p120 to p90 requires Rsp5 and the proteasome. (**A**) Domain organization of S. cerevisiae Mga2. The putative DNA binding domain (DBD) is shown in orange; the IPT (immunoglobulin-like fold, plexins, transcription factor) domain in blue; the ankyrin-repeat domain (Ank) in green; and the transmembrane segment (TM) in gray. The Rsp5 binding motif (LPKY) and the adjacent ubiquitination sites (Ub) are indicated. ( **B**) Flag-tagged full-length Mga2 (3xFlag-Mga2) was co-expressed with HA-tagged Rsp5 (HA-Rsp5) in Δpdr5 cells under the GAL1 promoter. Where indicated, the cells were incubated with Bortezomib. After solubilization in Triton, cell lysates were incubated with anti-Flag beads and the bound material was analyzed by SDS-PAGE and immunoblotting with antibodies to Flag (p120 and p90), HA (Rsp5), and ubiquitin (Ub). (**C**) As in (B), but comparing wild-type (WT) Mga2 with a mutant in which the Rsp5 binding mutant motif LPKY was changed to AAKA. An empty vector (EV) was used as control. (**D**) Flag-tagged full-length Mga2 was expressed from the TEF1 promoter. A cell lysate was separated into cytosol (p90) and membrane (p120) fractions. The membrane fraction was solubilized in Triton and both fractions were incubated with anti-Flag beads. Bound material was analyzed by SDS-PAGE and gel slices were treated with trypsin and subjected to mass spectrometry. Analysis of the peptide coverage (spectral counts) showed that p90 lacks peptides following residue 675. (**E**) AlphaFold3 predicted structure of the Ank domain. Residue 675 is located in a loop following the two N-terminal helices of the domain (teal).

We first tested whether the processing of Mga2’s p120 to the mature p90 fragment is dependent on the proteasome and Rsp5. Mga2 was tagged at the N-terminus with a FLAG epitope (FLAG-p120) and expressed under an inducible promoter in S. cerevisiae cells. The cells also expressed Rsp5 with a N-terminal hemagglutinin tag (HA-Rsp5) and lacked the drug transporter Pdr5 to enable the testing of proteasome inhibitors (*26*). The samples were subjected to immunoprecipitation (IP) with FLAG antibodies, followed by SDS-PAGE and blotting with antibodies to FLAG, HA, and ubiquitin. As expected, the addition of the proteasome inhibitor Bortezomib drastically reduced the conversion of p120 to p90 (**Fig. 1B**; upper panel; lane 4 versus 3). Rsp5 was co-precipitated under both conditions (middle panel), but ubiquitinated p120 accumulated only in the presence of the proteasome inhibitor (lower panel). To demonstrate that ubiquitination of p120 is mediated by Rsp5, we mutated the putative Rsp5-binding motif LPKY (**Fig. 1A**) to AAKA. The p120 mutant no longer bound Rsp5 and was not ubiquitinated (**Fig. 1C**; middle and lower panels; lane 6 versus 5). Rsp5-dependent ubiquitination of p120 was confirmed with purified proteins (**fig. S1D**): when bead-bound FLAG-p120 was incubated with purified Rsp5, the characteristic smear of ubiquitinated species was observed with wild-type p120 (lane 2 versus 1) but not the AAKA mutant (lane 6). Mutation of three lysine residues, previously implicated in the ubiquitination of p120 (K980, K983, K985)(*27*), to arginine (K3R mutant) reduced the modification, but did not abolish it (lane 4), indicating that other ubiquitination sites must exist.

To determine the C-terminal end of p90, i.e. the site at which the proteasome stops degrading p120, we overexpressed FLAG-p120 and purified the precursor p120 from the membrane fraction and the mature transcription factor p90 from the cytosolic fraction with anti-FLAG antibody beads. Both samples were subjected SDS-PAGE and excised bands were analyzed by mass spectrometry (**Fig. 1D**). For p120, we detected peptides spanning the entire sequence (top panel). For p90, however, we detected no C-terminal peptides after residue 675 (bottom panel), defining this position as the stoppage point of proteasomal degradation. This conclusion was confirmed by inserting the HA epitope at different positions of FLAG-p120 (**fig. S1E**): the HA epitope was detected in the mature p90 fragment if inserted at a position preceding residue 675 (residue 671 or 520), but not when inserted at downstream positions (residue 806 or C terminus). Residue 675 is located in a loop that follows the first two helices of the Ank domain (see scheme in **Fig. 1E**). Thus, the proteasome seems to have moved from the C-terminal ubiquitination sites through most of the folded Ank domain before it stopped, suggesting that not every folded domain can serve as a road block for proteasomal degradation. Interestingly, the remaining two Ank helices are not causing the stop of the proteasome either because deletion of the entire Ank domain (ΔAnk) still allowed the precursor to be processed into p90 (**fig. S1F**); the p90 band produced by the p120-ΔAnk mutant migrates at the same position as the one derived from wild-type (WT) p120 (p120-WT). A likely explanation is that proteasome movement is stalled at a defined distance from the IPT domain.

Although the proteasome normally leaves two helices of the Ank domain in the mature p90 protein, these are not required for the function of p90. A p90 fragment truncated at residue 626, which ends right after the IPT domain (see scheme in **Fig. 1A)**, fully rescued the growth of cells lacking both Mga2 and Spt23 in the absence of oleate (**fig. S1G**). Deletion of the Ank domain from full-length p120 also did not impair growth restoration in cells lacking Mga2 and Spt23 (**fig. S1G**), consistent with the observation that this domain is not required for the processing of p120 to p90 (**fig. S1F**). In agreement with previous reports (*12*), the processing to p90 is inhibited by unsaturated fatty acids (**fig. S1H**), but surprisingly, the Ank domain is not required for the inhibition (**fig. S1H**). The role of the Ank domain therefore remains to be clarified.

### The DBDs of Mga2 and p53 are related to one another

Both DALI (*28*) and Foldseek (*29*) searches with the AlphaFold-predicted structure (*30*) of Mga2’s DBD returned as the top hit the DBD of the mammalian transcription factor p53. p53 is a crucial tumor suppressor protein, whose primary function is to protect the body from cancer by regulating cell division and ensuring that damaged cells cannot multiply (*31*). p53 pauses the cell cycle, initiates DNA repair, and triggers apoptosis if the DNA damage is too severe to be fixed. Although p53 is widespread in metazoans, it has been thought to be absent in fungi and other lower organisms. However, the superposition of the DBDs of Mga2 and p53 shows that the cores of the domains are highly related (**Fig. 2A**). Some peripheral regions are different. For example, the p53 helix that binds DNA is lacking in Mga2, and instead a different helix is predicted by AlphaFold3 to contact the DNA (**Fig. 2, A to C**). Three basic residues in Mga2 (R252, R253, K254) are expected to bind into the major groove of double-stranded DNA (Fig. 2c). The residues in the putative DNA binding helix of Mga2 and Spt23, including the three basic residues, are evolutionarily conserved as analyzed by Consurf (*32*) (**fig. S2A**).

**Fig. 2.**
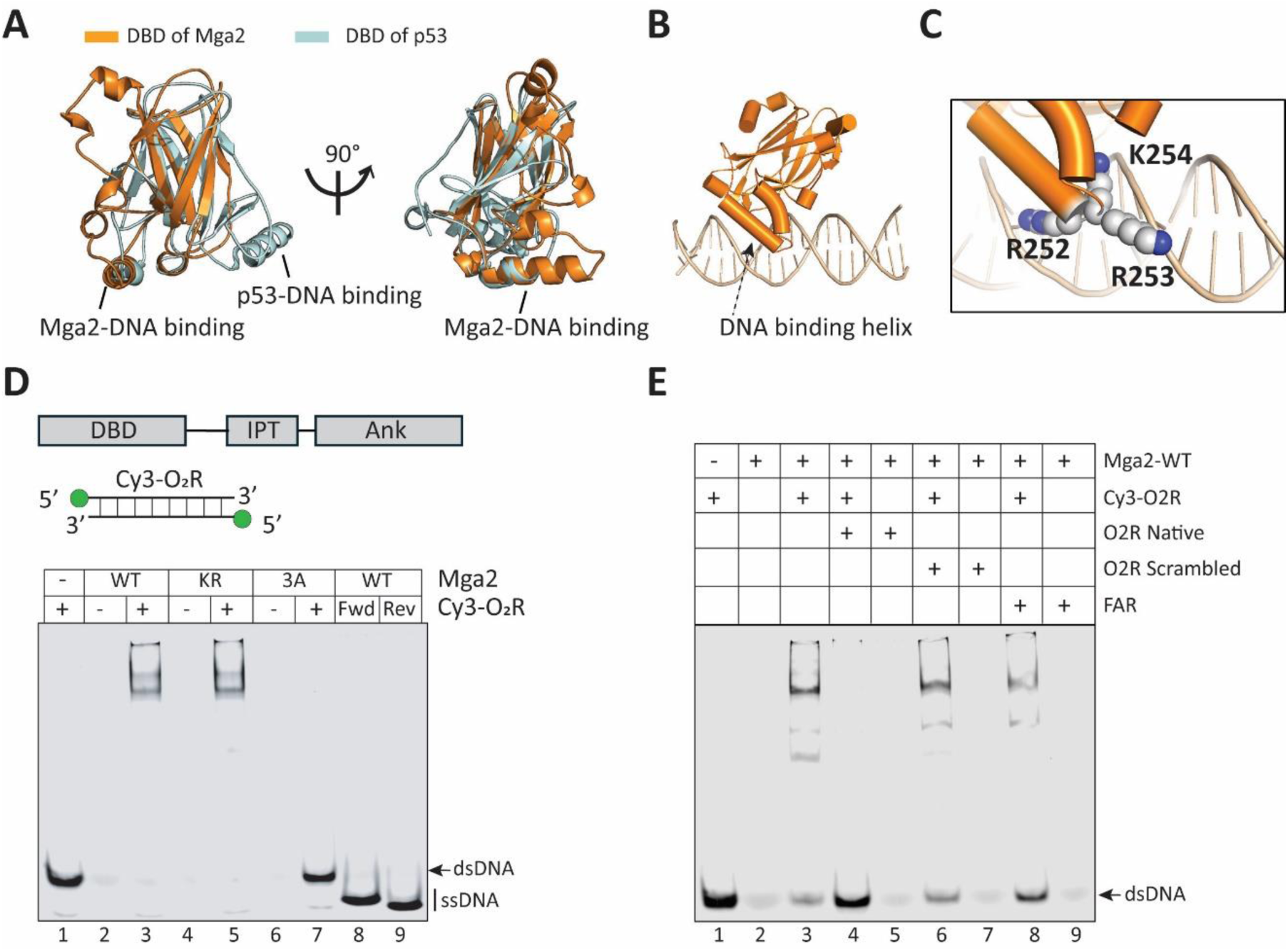
Mga2’s DBD directly binds to DNA. (**A**) Superposition of the DBD structure of p53 (grey; PDB code: 3KMD) with the DBD structure of Mga2 (orange) predicted by AlphaFold3 (r.m.s.d: 2.5Å). Shown are two different views of cartoon models. (**B**) AlphaFold prediction of Mga2’s DBD in complex with the double-stranded O2R promoter sequence (50bp). (**C**) Magnified view of the interaction with DNA. Three conserved positively charged residues (R252, R253, K254) bind into the major groove and are shown as spheres. (**D**) Purified Mga2 constructs spanning the region from the DBD to the Ank domain (see scheme) were incubated with double-stranded (ds) O2R promoter DNA fluorescently labeled with Cy3 (Cy3-O2R). The constructs contained either wild-type (WT) DBD or mutations in the DNA-binding helix (KR and 3A). All samples were subjected to native gel electrophoresis and analyzed by fluorescence scanning. Single-strand (ss) O2R sequences (5’-3’: Fwd, 3’-5’: Rev) were used as controls. (**E**) As in (D), but the WT construct was incubated with Cy3-O2R and an excess of unlabeled competitor, as indicated.

To test the predicted interaction between the DBD and DNA, we performed gel-shift assays with constructs that contain either wild-type DBD or mutants in which either all three basic residues (R252, R253, K254) are mutated to alanine (3A mutant) or K254 is mutated to arginine (KR mutant). Mga2 fragments spanning the region from the DBD to the Ank domain were expressed as fusions with the maltose-binding protein (MBP) in E. coli and purified on an amylose resin, followed by chromatography on a heparin column to remove non-specifically bound bacterial DNA (**figs. S2, B and C**). The purified proteins were incubated with a fluorescently labeled double-stranded (ds) DNA fragment corresponding to the O2R promoter region, and subjected to native gel electrophoresis (**Fig. 2D**). Wild-type (WT) protein and the KR mutant shifted the fluorescent dsDNA probe to a high-molecular weight position (lanes 3 and 5), whereas the 3A mutant did not (lane 7). No binding of the WT protein was seen with the individual, single-stranded DNAs in the probe (lanes 8 and 9). The interaction was specific for the O2R promoter sequence, as it was prevented by the addition of an excess of unlabeled native, but not scrambled, DNA (**Fig. 2E**; lanes 3 versus 4 and 6). Mga2 protein also bound to the FAR promoter DNA sequence, albeit weaker than to the O2R sequence (lane 8). No binding was observed with a purified MBP-fusion of the DBD domain alone (**fig. S2, D to F**), perhaps because efficient binding requires dimerization through the IPT domain.

### DNA binding-dependent degradation of p90 in the nucleus

Consistent with the in vitro DNA-binding results, the overexpression of the WT fragment 1-626 (called p90 from here on) or of the corresponding KR mutant of Mga2 rescued the growth of cells lacking both Mga2 and Spt23, whereas the 3A mutant was inactive (**Fig. 3A**). Full-length Mga2 containing the 3A mutations also did not rescue the growth of the double-deletion mutant (**fig. S3A**). As expected, the Rsp5-binding mutant (AAKA mutant) was also inactive, whereas ablation of the known ubiquitination sites (K3R mutant) had no effect (**fig. S3A**), consistent with the existence of additional, unidentified ubiquitination sites.

**Fig. 3.**
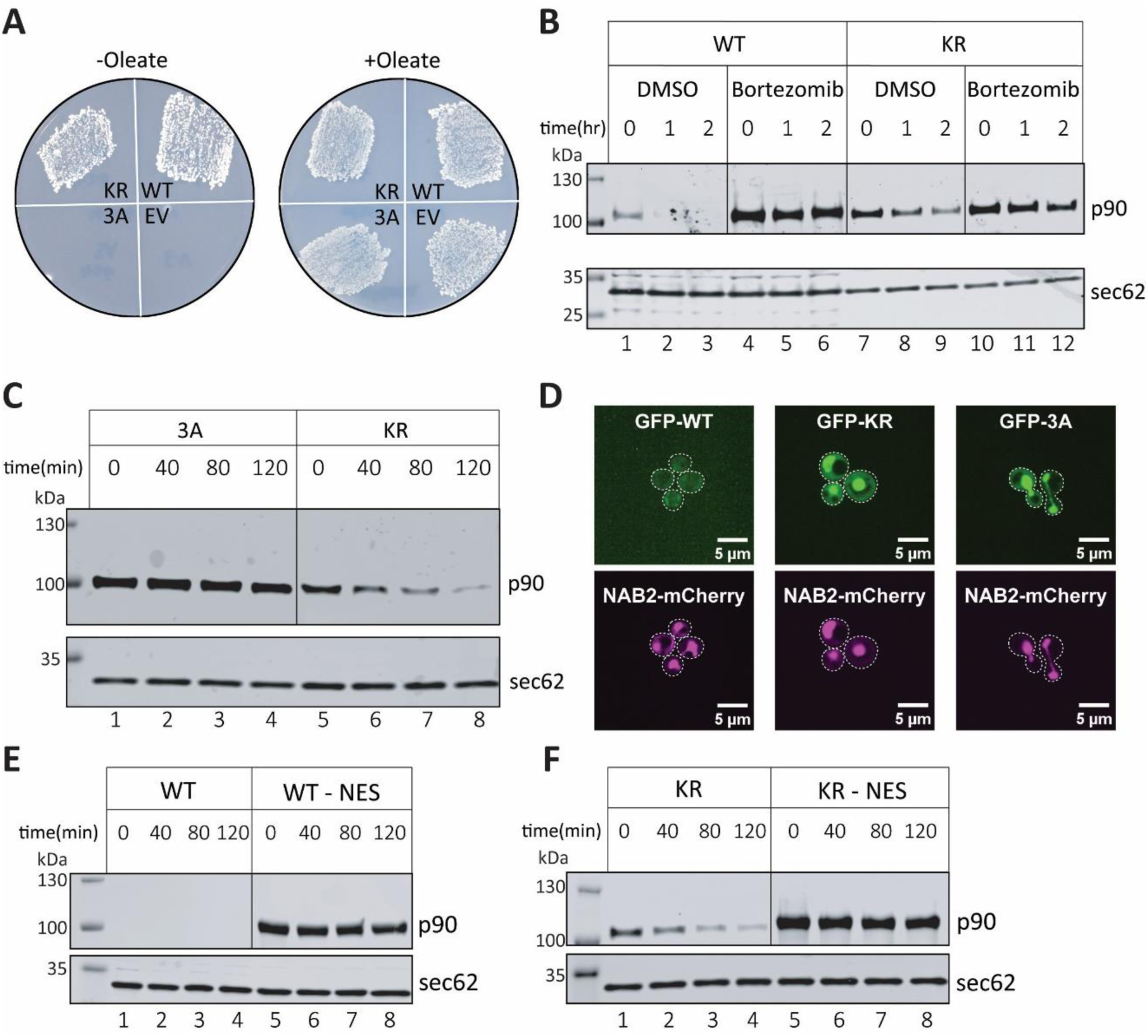
Nuclear degradation of p90 depends on DNA binding. (A) Flag-tagged wild-type (WT) p90 (Flag-p90; residues 1-626) or the corresponding KR and 3A mutants were over-expressed in cells lacking both Mga2 and Spt23. An empty vector (EV) was used as control. The cells were grown on plates lacking or containing oleate at 30°C for three days. (B) HA-tagged p90 wild-type (WT) p90 (2xHA-p90; residues 1-626) or the corresponding KR mutant were over-expressed in Dpdr5 cells. The cells were treated with DMSO or Bortezomib for two hours, before adding cycloheximide. Samples were taken at different times points and cell lysates were analyzed by SDS-PAGE and immunoblotting with HA antibodies. Blotting for Sec62 served as a loading control. (**C**) Flag-p90 carrying the KR or 3A mutations was over-expressed in Dmga2 cells. Cycloheximide chase experiments were performed as in b, except that immunoblotting was performed with Flag antibodies. (**D**) GFP-tagged wild-type p90 (GFP-WT; residues 1-626) or the corresponding KR and 3A mutants were over-expressed in cells stably expressing the nuclear marker NAB2-mCherry. The cells were analyzed for GFP and mCherry fluorescence by confocal microscopy. The boundaries of the cells are indicated by broken lines. (**E**) HA-tagged wild-type p90 (residues 1-626) or p90 fused at the C-terminus to two consecutive nuclear export signals (LALKLAGLDI) (WT-NES) were over-expressed in Δmga2 cells. Cycloheximide-chase experiments were performed as in (B). (**F**) As in E, but with the KR mutant.

The WT p90 fragment was efficiently degraded in yeast cells by the proteasome, as shown by the effect of the Bortezomib in cycloheximide-chase experiments (**Fig. 3B**; lanes 1-3 versus 4-6). The KR mutant was considerably more stable, but still a substrate of the proteasome (lanes 7-12). In contrast, the 3A mutant was completely stable (**Fig. 3C**; lanes 1-4 versus 5-8). Consistent with these results, WT p90 fused at the N-terminus to GFP (GFP-WT) could barely be detected in cells, whereas the corresponding fusions of the KR and 3A mutants were visible inside the nucleus (**Fig. 3D**). A GFP fusion of the full-length 3A mutant also localized to the nucleus (**fig. S3B**). These results confirm that p90 moves into the nucleus and indicate that DNA binding is required for the degradation of p90. p90 degradation likely occurs inside the nucleus because the fusion of WT p90 or the KR mutant to a nuclear export signal (NES)(*33*) rendered the proteins stable in cycloheximide-chase experiments (**Fig. 3, E and F**; lanes 5-8 versus 1-4). Furthermore, the deletion of the p90 region 335-526, which encompasses putative nuclear import signals, impaired the accumulation of the protein in the nucleus (**fig. S3C**), stabilized the KR mutant protein (**fig. S3D**), and compromised the ability of the KR mutant to promote Ole1 transcription (**fig. S3E**).

### DNA binding in vivo through the DBD

Next, we tested DNA binding in vivo. We expressed HA-tagged p90 carrying the KR or 3A mutations in cells lacking Mga2, and induced crosslinks to DNA with formaldehyde. After shearing the DNA and immunoprecipitation with HA antibodies, the crosslinks were reversed, and the co-precipitated DNA was analyzed by sequencing. The KR mutant bound preferentially to Ole1’s promoter region of 750 bp that contains both the FAR and O2R sequences (**Fig. 4A**). As expected, less binding was seen with the 3A mutant. These results were confirmed by analyzing the O2R promoter region separately with qPCR (**Fig. 4B**). DNA binding could not be analyzed with the WT protein, as it is very unstable and thus present at low concentrations.

**Fig. 4.**
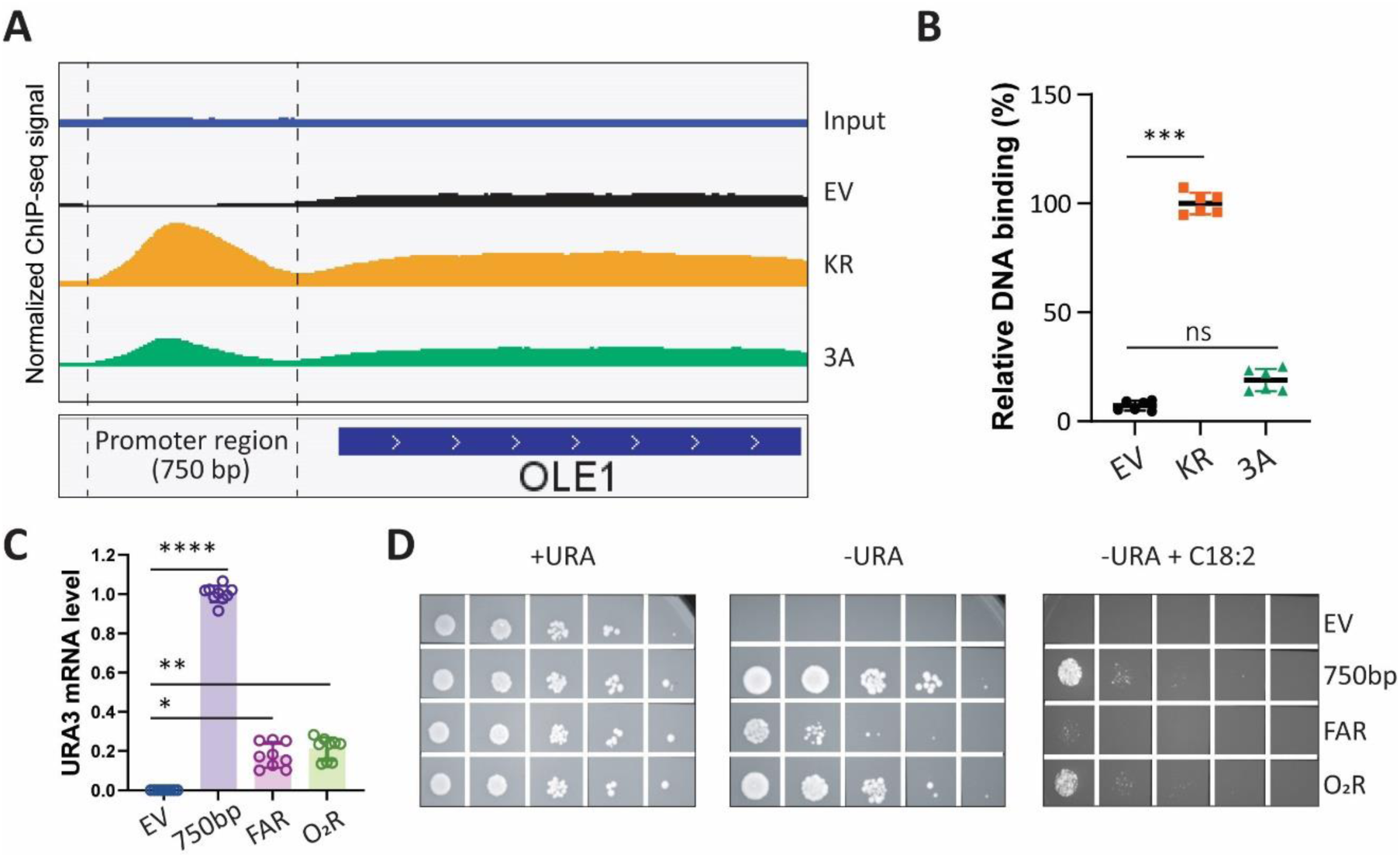
In vivo DNA binding of p90. (A) HA-tagged p90 (2xHA-p90; residues 1-626) bearing the KR or 3A mutations was over-expressed in Dmga2 cells. A control was performed with an empty vector (EV). The cells were treated with formaldehyde, and a cell lysate was sonicated to shear the DNA. After immunoprecipitation with HA antibodies, the crosslinks were reversed, and the precipitated DNA was subjected to next-generation sequencing. Normalized read coverage across the OLE1 locus is shown. A sample was analyzed without immunoprecipitation (Input). The promoter and coding regions of the Ole1 gene are indicated. (**B**) The DNA samples in a were analyzed by qPCR with primers flanking the O2R promoter region (ChIP-qPCR). DNA binding is shown relative to that of the KR mutant. (**C**) The entire promoter region of the Ole1 gene (750 bp), or the FAR (100 bp) or O2R (50 bp) regions, were fused to the URA3 gene and expressed from a centromeric plasmid in Dura3 cells. An empty centromeric plasmid was used as control (EV). The RNA was extracted from cells and Ura3 expression quantified by RT-qPCR. URA3 mRNA levels are expressed relative to that with the entire promoter region. (**D**)The cells in c were serially diluted and spotted onto plates containing or lacking uracil (+URA and -URA, respectively). The plates on the right lacked uracil and contained linoleic acid (C18:2). All plates were incubated at 30°C for three days.

We used fusions of the Ole1 promoter region with the URA3 gene to test the importance of the FAR and O2R sequences. A region of 750bp that contains both elements caused robust transcription of the URA3 gene, whereas each element alone was much less active (**Fig. 4C**). Both the 750 bp region and the O2R element allowed cells to grow on uracil-lacking medium, whereas the FAR element was less effective (**Fig. 4D**). Thus, in agreement with our in vitro results (**Fig. 2E**), the O2R element causes stronger p90 binding. The expression of all promoter fusions was inhibited when the fatty acid C18:2 was added, indicating that the unsaturated fatty acid regulation of Ole1 expression is mediated by both the FAR and O2R elements.

### Mga2 has a p53-like trans-activation domain (TAD)

We noticed that Mga2 has a sequence near its N-terminus that resembles the trans-activation domain 1 (TAD1) of p53 (**Fig. 5A**). TAD1 is required for the degradation of p53, a process triggered by the ubiquitin ligase Mdm2 (*34*). Mdm2 is the chief regulator of p53, as it maintains p53 at low baseline levels in unstressed cells, preventing unwanted cell cycle arrest or apoptosis. TAD1 binds to a domain of Mdm2 through a helix whose hydrophobic amino acid residues bind into a groove of this domain (*35*) (**Fig. 5B**).

**Fig. 5.**
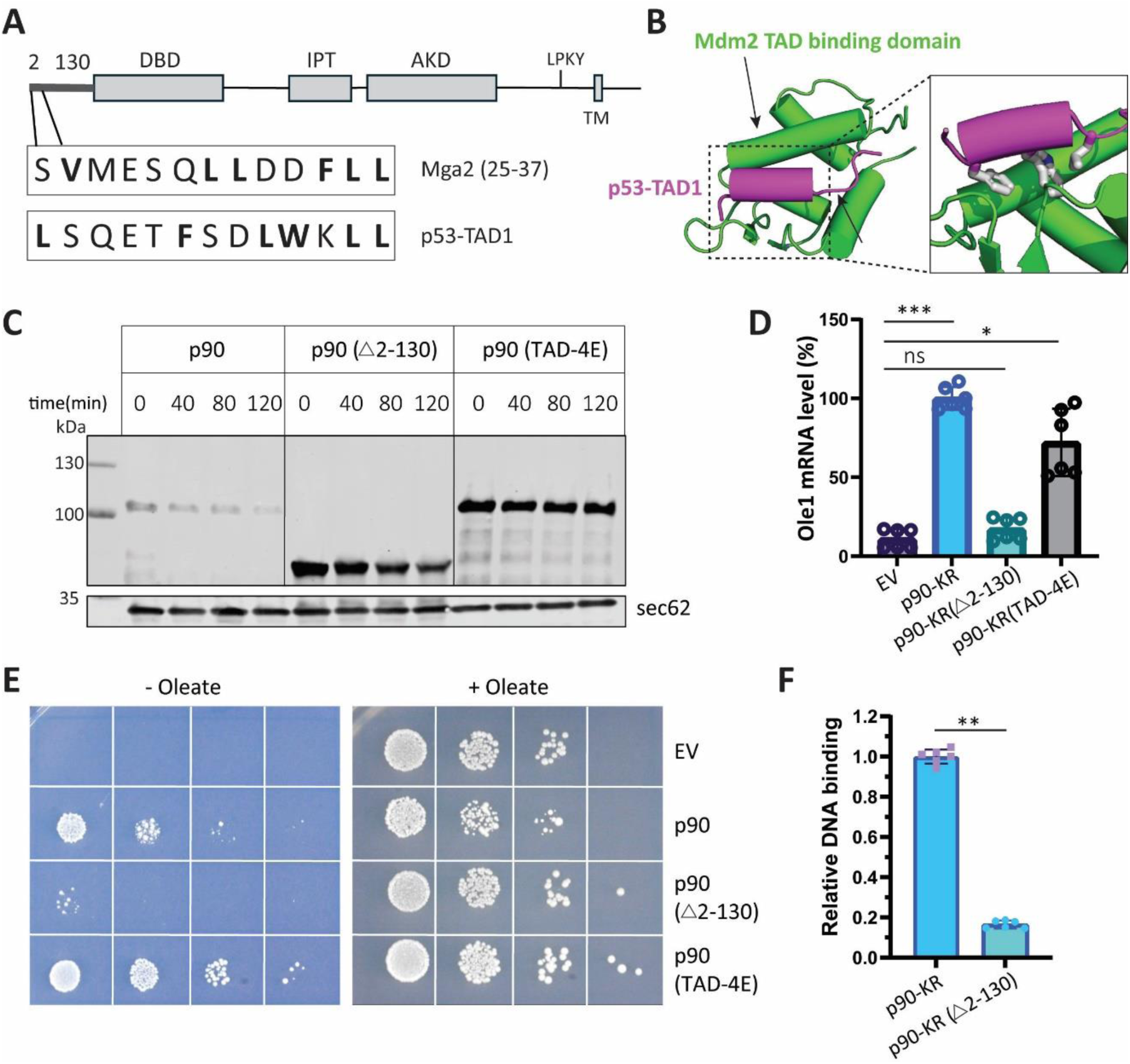
Mga2 contains a trans-activation-like domain required for p90 degradation. (**A**) Comparison of the Mga2 amino acid sequence 25-37 with the trans-activation domain 1 (TAD1) of p53. Hydrophobic residues are shown in bold. (**B**) Cartoon model of a crystal structure of p53-TAD1 bound to the TAD-binding domain of Mdm2 (PDB: 4HFZ). TAD1 and Mdm2 are shown in magenta and green, respectively. The hydrophobic residues of TAD1 are shown as sticks in the magnified view. (**C**) HA-tagged p90 wild-type (WT) p90 (2xHA-p90; residues 1-626) or the indicated mutants were over-expressed in Δmga2 cells. Cycloheximide was added and samples were taken at different time points. Cell lysates were subjected to SDS-PAGE and analyzed by immunoblotting with HA antibodies. Blotting for Sec62 served as a loading control. (**D**) 2xHA-p90 containing the KR mutation (p90-KR) and variants that additionally have amino acids 2-130 deleted (p90-KR (Δ2-130)) or bear mutations in the TAD-like sequence (p90-KR(TAD-4E)) were over-expressed in Δmga2 cells. As a control, cells were transformed with an empty vector (EV). The RNA was extracted and Ole1 mRNA levels determined by RT-qPCR. The levels of Ole1 transcripts are expressed relative to those induced by p90-KR. (**E**) Wild-type p90 (residues 1-626) or mutants (p90(Δ2-130) or p90-(TAD-4E)) were expressed from a centromeric plasmid in cells lacking both Mga2 and Spt23. An empty vector (EV) was used as control. The cells were incubated at 30°C for three days on plates lacking or containing oleate. (**F**) HA-tagged p90-KR or p90-KR (Δ2-130) were over-expressed in Δmga2 cells. Binding to the O2R promoter of the Ole1 gene was determined by ChIP-qPCR as in Fig. 4B.

Like the TAD1 of p53, the TAD-like sequence of Mda2 contains hydrophobic and acidic amino acids (**Fig. 5A**). Yeast does not have a Mdm2 homolog but, as in the case of p53, the TAD-like sequence of Mga2 causes degradation of the transcription factor: mutation of four hydrophobic amino acids in the TAD-like sequence to glutamates (TAD-4E mutant) completely stabilized p90 (**Fig. 5C**). To test the effect of p90 and its mutants on Ole1 expression, we introduced the stabilizing KR mutation into p90, which resulted in more robust transcription. Unexpectedly, the TAD-4E/KR mutant allowed considerable transcription of the Ole1 gene (**Fig. 5D**). Furthermore, p90 carrying only the 4E mutations permitted the growth of cells lacking endogenous Mga2 and Spt23 (**Fig. 5E**). Thus, the 4E mutant of p90 can still trigger Ole1 transcription, suggesting that the TAD-like sequence of Mga2 is not required for transcription. Mga2 seems to have other sequences that function as actual TADs, as deleting a larger N-terminal unstructured region (Δ2-130) from the KR mutant drastically reduced Ole1 transcription (**Fig. 5D**). The Δ2-130/KR mutant also bound much weaker to the O2R promoter than the intact p90/KR protein (**Fig. 5F**). Furthermore, deleting the 2-130 region from the wild-type protein prevented the growth of cells lacking Mga2 and Spt23 (**Fig. 5E**) without affecting nuclear import (**fig. S3C**). This Δ2-130 mutant was also significantly more stable than the wild-type protein (**Fig. 5C**). Taken together, these results show that p90 has a TAD-like sequence that causes its degradation, and probably additional TADs that are required for Ole1 transcription.

### Biological significance of p90 degradation

The degradation of p90 was insensitive to the addition of unsaturated fatty acids or CoCl_2_, the presence of which mimics low oxygen conditions (*5*) (**Fig. 6A**). Thus, the physiological regulation of Ole1 expression seems to occur exclusively by changes in the synthesis of p90. However, as in the case of p53, rapid degradation of the mature transcription factor might still be physiologically important, because it would keep Ole1 expression low under normal conditions. To investigate the biological consequence of impaired p90 degradation, we used the KR mutant of p90. This mutant is considerably more stable than the wild-type protein and therefore has a higher steady-state level (**Fig. 6B**; top panel), but it shows equal DNA binding in vitro (**Fig. 2D**). Consistent with these properties, the KR mutant generated more Ole1 transcripts in Mga2-lacking cells than the wild-type protein (**Fig. 6B**; bottom panel). The DNA binding-defective 3A mutant was present at even higher levels than the KR mutant (**Fig. 6B**, top panel), but was totally inactive in promoting Ole1 transcription (**Fig. 6B**; bottom panel). Interestingly, the transcription level of Ole1 correlated with the number of lipid droplets, as shown by microscopy after staining yeast cells with BODIPY 493/503 (**Fig. 6C**) or by analyzing BODIPY 493/503-labeled cells with FACS (**Fig. 6D**); the KR mutant generated considerably more lipid droplets than WT p90, whereas the 3A mutant was inactive. These results suggest that unsaturated acyl chains generated in excess by Ole1 are stored as triglycerides in lipid droplets. Indeed, analysis of the fatty acyl chains by gas chromatography and mass spectrometry showed that the ratio of unsaturated to saturated acyl chain is higher in cells expressing the KR mutant than in cells expressing WT p90 or the 3A mutant (**Fig. 6E**). Lipid droplet formation is essential for cell viability if Ole1 is upregulated, as the expression of WT p90 or the KR mutant in cells lacking the four enzymes required for lipid droplet formation (Dga1, Lro1, Are1, and Are2)(*36*) caused lethality, in contrast to the expression of the 3A mutant (**Fig. 6F**). Taken together, these results indicate that efficient p90 degradation and consequent moderation of Ole1 expression as well as the formation of lipid droplets are both required to prevent the accumulation of toxic unsaturated fatty acids.

**Fig. 6.**
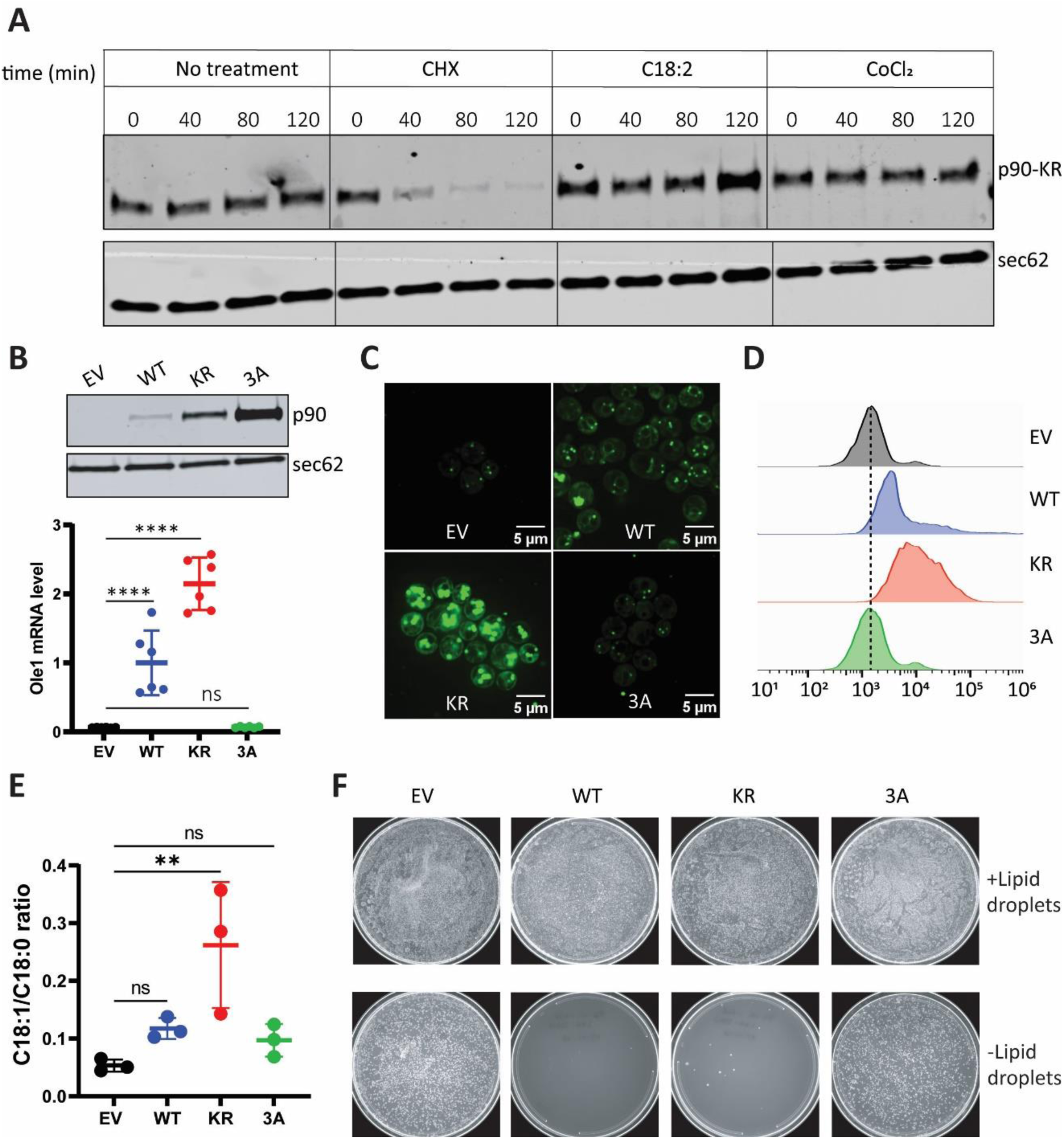
Impaired p90 degradation induces Ole1 transcription and lipid droplet formation. (**A**) HA-tagged p90-KR (residue 1-626) was over-expressed in Δmga2 cells. The cells were incubated for the indicated times without additions (no treatment) or with cycloheximide, linoleic acid (C18:2; 200 μM) or CoCl_2_ (3 mM). Total cell lysates were subjected to SDS-PAGE and immunoblotting with HA antibodies. Blotting for Sec62 served as a loading control. ( **B**) Flag-tagged wild-type (WT) p90 (residues 1-626) or the corresponding KR and 3A mutants were expressed from the TEF1 promoter on a CEN plasmid in Δmga2 cells. As a control, cells were transformed with an empty vector (EV). Cell lysates were subjected to SDS-PAGE and immunoblotting with Flag antibodies (bottom panel). Ole1 transcript levels were measured by RT-qPCR (top panel). The levels of Ole1 transcripts are expressed relative to those induced by WT p90. ( **C**) The indicated cells were stained with BODIPY 493/503 and visualized by confocal microscopy. ( **D**) As in (C), but BODIPY 493/503-stained cells were analyzed by FACS. (**E**) Lipids were extracted from the indicated cells. The acyl chains were converted to fatty acid methyl esters and analyzed by gas chromatography and mass spectrometry (GC-MS). ns, not significant; **, p=0.005. (**F**) Wild-type (WT) p90 or the KR or 3A mutants were expressed from the TEF1 promoter on a CEN plasmid in either Δmga2 cells (+Lipid droplets) or Δmga2 cells lacking the four enzymes required for lipid droplet formation (Dga1, lro1, Are1, Are2; -Lipid droplets). The cells were incubated on plates at 30°C for three days.

## DISCUSSION

We have characterized the molecular pathway by which the transcription factor Mga2 controls the expression of the fatty acid desaturase Ole1 and thereby maintains lipid homeostasis. The ER membrane-anchored p120 precursor is first ubiquitinated by the E3 ligase Rsp5 (**Fig. 7, step 1**) and then degraded by the 26S proteasome to generate the mature transcription factor p90 (**Fig. 7, step 2**). Degradation of p120 is likely initiated through insertion of a polypeptide loop into the proteasome (step 2). This loop must be located close to the TM segment, perhaps in proximity of the previously identified ubiquitination sites (*27*). The proteasome then proceeds towards the N-terminus and processively degrades the precursor until its movement is stopped. Although in vitro studies have established that tightly folded domains can arrest proteasomal degradation (*37–39*), our results show that the proteasome traverses the folded Ank domain before stalling at a fixed distance from the IPT domain. This implicates the IPT domain as a specific degradation-stop signal for p120 processing, analogous to the proposed role of the IPT domain in NF-κB processing. IPT domains are found in several other transcription factors, and it will be interesting to determine whether partial proteasomal processing is a general feature of this protein family.

**Fig. 7.**
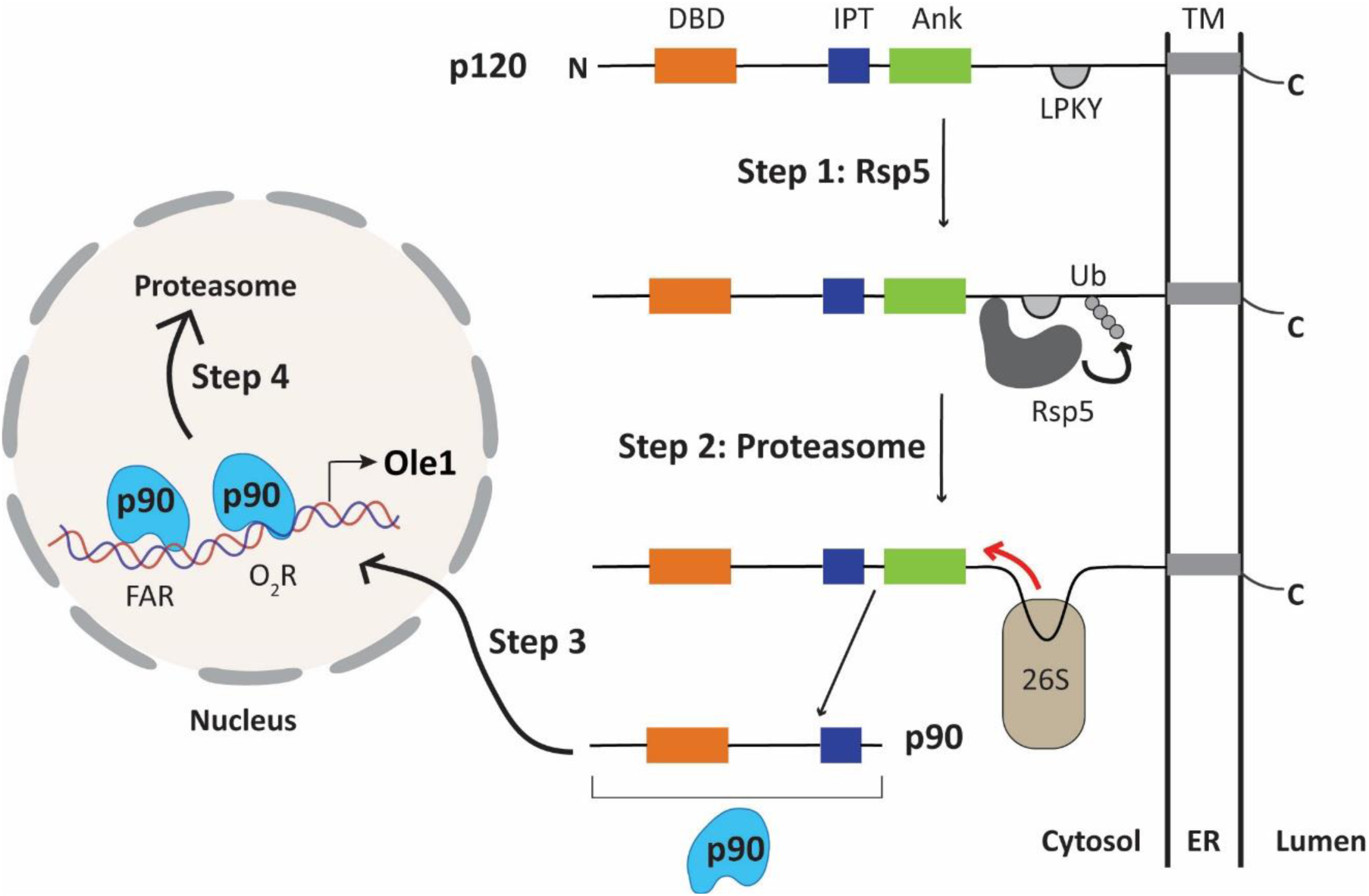
Model of the Mga2 pathway. Step 1: The ER-bound p120 precursor binds the ubiquitin ligase Rsp5 through its LPKY motif and is ubiquitinated. Step 2: A loop of p120 inserts into the 26S proteasome. The proteasome moves towards the N-terminus and degrades p120 until it is stalled near the IPT domain, resulting in cytosolic p90. Step 3: p90 moves into the nucleus, binds to the FAR and O2R promoter elements of the OLE1 gene, and activates transcription. Step 4: Once bound to DNA, p90 is degraded by the proteasome.

The mature p90 fragment translocates to the nucleus to activate Ole1 transcription (**Fig. 7, step 3**). Strikingly, the DNA-binding domain and a trans-activation-like segment of p90 are structurally and functionally homologous to those of the mammalian tumor suppressor p53. The DBD binds specifically to two promoter elements in the Ole1 gene — the O2R and FAR regions — with higher affinity for O2R, despite the similarity between the two elements was found by pairwise alignment (*40*). Although these elements were originally characterized as mediating distinct regulatory inputs (*41*), our findings raise the possibility that they cooperate in controlling Ole1 transcription. How the DBD achieves sequence-specific DNA recognition remains an open question, as AlphaFold modeling predicts binding within the major groove regardless of sequence context.

The trans-activation-like segment of p90 (aa 25-37) is related to TAD1 of p53. Both are required for the proteasomal degradation of the mature transcription factors. In p53, TAD1 recruits the E3 ligase Mdm2 to drive polyubiquitination and subsequent degradation. The mechanism by which the TAD-like sequence of Mga2 triggers p90 degradation is unknown, as no cognate ubiquitin ligase has yet been identified. As with p53, p90 degradation is initiated inside the nucleus and depends on DNA binding and likely active transcription (**Fig. 7, step 4**). Importantly, transcription does not require p90 degradation, as our TAD-4E mutant — which is completely stable — retains much of its transcriptional activity. How DNA binding and transcription of the Ole1 gene are mechanistically coupled to p90 degradation remains an important open question.

The TAD1 motif of p53 not only binds Mdm2, but also p300, a histone acetyltransferase involved in chromatin remodeling (*34*). Intriguingly, the TAD1-binding domain of Mdm2 has homologs in other chromatin-remodeling complexes. This domain can structurally be aligned with domains of BAF60 and Swp73, components of the mammalian and yeast SWI/SNF complexes, respectively, as well as with a domain of Rsc6, a component of the yeast RSC complex (**fig. S4A**). AlphaFold predicts that the TAD1-related sequence of Mga2 binds to Swp73 and Rsc6 in the same way as TAD1 of p53 to Mdm2 (**fig. S4B**), suggesting this sequence may also serve to recruit chromatin-remodeling complexes. However, our results show that the motif is dispensable for activating Ole1 transcription. Consistent with our evidence that other TAD sequences activate the transcription of Ole1, AlphaFold predicts that an adjacent sequence also interacts with the domains of Swp73 and Rsc6 (fig. S4c).

Like p53 and many other transcription factors (*42*), p90 is rapidly degraded following target gene activation, a mechanism that likely enables cells to rapidly respond to changing conditions. p90 turnover does not appear to be regulated by the physiological signals that control Ole1 expression —unsaturated fatty acids or oxygen availability, but is important to keep Ole1 transcription at a low basal level. Compromising p90 turnover causes excessive accumulation of unsaturated fatty acids, which are then stored as triglycerides in lipid droplets to prevent lipotoxicity. This reveals an additional layer of regulation: unsaturated fatty acids not only suppress p90 generation by inhibiting p120 processing, but are also detoxified by conversion into neutral lipids when they accumulate in excess.

Our results reveal a striking and unexpected similarity between Mga2 and p53 at the levels of structure, mechanism, and regulatory logic, despite their entirely different biological roles. Although we begin to understand the Mga2 pathway, many aspects remain to be clarified, such as how exactly proteasome degradation of the p120 precursor is stalled, how the p90 fragment is degraded in the nucleus, and how the pathway is regulated by unsaturated fatty acids.

## MATERIALS AND METHODS

### Co-immunoprecipitation of Mga2 and Rsp5

3xFlag-Mga2 or the corresponding Flag-tagged Rsp5-binding mutant were cloned into the pBEVY-GU vector. HA-Rsp5 was cloned into the pBEVY-GL vector. Δpdr5 cells (*43*) were transformed with these plasmids and grown on CSM-Leu-Ura plates. The cells were cultured in CSM-Leu-Ura medium containing 2% raffinose and 0.1% glucose overnight at 30°C before harvesting (total OD600 40-60). The cells were washed and incubated for 4 hours in CSM-Leu-Ura medium containing 2% galactose to induce the expression of Mga2 and Rsp5. DMSO or 80 μM of bortezomib (MCE, HY-10227) in DMSO were added together with galactose. The cells were resuspended in lysis buffer (50 mM HEPES, 100 mM Nacl, 10% glycerol, pH7.4, 2 x protease inhibitor cocktail (MCE.HY-K0010), 1 mM 1,10-phenanthroline, 80 μM bortezomib, 1mM PMSF) and lysed by bead-beating. After centrifugation at 950xg for 30s, 1% Triton X-100 was added to the supernatant for 1 hr at 4°C. The samples were centrifuged at 18,000xg for 10 min and the supernatant incubated with magnetic anti-Flag M2 resin (Sigma, M8823) at 4°C for 1 hr. The resin was washed five times with lysis buffer containing 0.1% Triton X-100. Bound material was eluted with 40 or 50 μl of 1xSDS loading buffer.

### Purification of p120 and p90

Flag-tagged full-length Mga2 (Flag-Mga2) was cloned into the pRS416 vector behind the TEF1 promoter, and the plasmid was transformed into Δmga2 cells (horizon; clone ID 5968). The cells were grown in 800ml to 1L of CSM-Ura medium, sedimented, and resuspended in lysis buffer (50 mM HEPES, 400 mM Nacl, 5% glycerol, pH7.4, 1 mM TCEP, 2 x protease inhibitor cocktail, 1mM PMSF). Lysis was performed by bead-beating in multiple tubes. A crude cell lysate was harvested by centrifugation at 950xg for 30s, and further centrifugation at 8,000xg for 5 min. The supernatant was separated into cytosol and membrane fractions by centrifugation at 100,000xg for 45 min. The cytosol fraction was incubated with anti-Flag M2 resin (Sigma, A2220) at room temperature (RT) for 1 hr and washed five times with 10 ml lysis buffer containing 0.02% dodecyl-β-D-maltoside (DDM). The bound material was eluted with 20ul of 4xSDS loading buffer. The membrane pellet was resuspended in 1 ml of lysis buffer and solubilized with 1% Triton X-100 at 4oC for 1.5 hr. After centrifugation at 18,000xg for 10 min, the supernatant was incubated with anti-Flag M2 resin at 4oC for 1.5 hr. The beads were washed five times with 10 ml lysis buffer containing 0.1% Triton X-100. The bound material was eluted with 20ul of 4xSDS loading buffer. All samples were subjected to SDS-PAGE and staining with Coomassie blue G-250, and the relevant bands were excised and subjected to mass spectrometry.

### Mass spectrometry analysis of p120 and p90

The excised bands were treated with trypsin and the gel-extracted peptides (*44*) were desalted with StageTip, dried by vacuum centrifugation, and reconstituted in 5% acetonitrile, 5% formic acid for LC-MS/MS processing.

Mass spectrometry data were collected using an Orbitrap Astral mass spectrometer (Thermo Fisher Scientific, San Jose, CA) coupled with a Neo Vanquish liquid chromatograph. Peptides were separated on a 110 cm µPAC C18 column (Thermo Fisher Scientific). For each analysis, ∼0.5 μg were loaded onto the column. The peptides were separated using a 60 min gradient of 5 to 29% acetonitrile in 0.125% formic acid with a flow rate of 300 μL/min.

The scan sequence began with an Orbitrap MS1 spectrum using the following parameters: resolution 60,000, scan range 350−1350 Th, automatic gain control (AGC) target 200%, maximum injection time 50ms, RF lens setting 50%, and centroid spectrum data type. FAIMS was enabled with using Top30 setting for each. Samples were analyzed three times with two compensation voltages (CV) sets: −30V, −40V, −45V, −50V, and −60V for two replicate analyses and one with −25V, −35V, −45V, −55V, −60. Astral data acquisition included AGC 200%, maximum injection time 15ms, isolation window 1.2 Th, normalized collision energy (NCE) 27%, and centroid spectrum data type. In addition, unassigned and singly charged species were excluded from MS2 analysis and dynamic exclusion was set to 15 s across CVs.

Mass spectra were processed using a Comet-based in-house software pipeline. MS spectra were converted to mzXML using MSconvert. Database searching included the protein of interest (Mga2) plus common contaminants which was concatenated with a reverse database composed of all these protein sequences in reversed order. The digest was set to non-specific. Searches were performed using a 3 Da precursor ion tolerance. Product ion tolerance was set to 0.02 Th. Oxidation of methionine residues (+15.9949 Da) was set as a variable modification. PSM filtering was performed manually considering only tryptic peptides, an XCorr>1.5, and an absolute value of PPM mass tolerance <10. Spectra were manually validated.

### CRISPR-Cas9 editing of Mga2 and Spt23

To attach 2xHA tags at the N terminus of Mga2 or Spt23, the pML104 plasmid (Addgene #67638) was digested with SwaI and BclI. Guide RNA sequences (5’-3’: GCAGAACAGTGAGTTCTTAA for Mga2; 5’-3’: CTGAAAATGATGAGTGGCAC for Spt23) were ligated into the pML104 plasmid using an overnight incubation at 16oC with T4 ligase. The pML104 plasmids together with donor sequences (see supplementary Table 1) were transformed into the BY4741 yeast, and cells were selected on CSM-Ura plates. Single colonies were grown in YPD medium, and 500 ul of cells were incubated with 20 mM NaOH at 100°C for 10 min to extract the gDNA. Diagnostic PCR was performed (Mga2_check_F&R for Mga2; Spt23_check_F&R for Spt23) and the amplicon was sequenced to confirm successful knock-in. For the deletion of the Ank domain of Mga 2(ΔAnk), the guide RNA sequence (5’-3’: ACAGTAATTATTCTATTAGC) was similarly inserted into the pML104 plasmid. The pML104 plasmid together with the donor sequence (see supplementary Table 1) was transformed into 2xHA-Mga2 yeast cells. Colonies on CSM-Ura plates were analyzed by diagnostic PCR (Mga2_check_F&R) and sequencing. 2xHA-Mga2 WT or ΔAnk, and 2xHA-Spt23 strains were grown in YPD medium, and immunoprecipitation was performed with anti-HA resin to probe for p120 and p90. For experiments involving unsaturated fatty acid, 200 μM of C18:2 (Sigma, L1376; prepared as 1% stock solution in 10% IGEPAL-630) was added to yeast cell cultures at an OD of 0.30. The cells were further cultured at 30oC for 4 hours and harvested for lysis.

### Immunoprecipitation of p120 and p90

Typically, 40-80 OD cells were used for detecting 2xHA-tagged Mga2 expressed from the chromosome or centromeric plasmids. The cells were resuspended in lysis buffer (50 mM HEPES, 400 mM Nacl, 5% glycerol, pH7.4, 1 mM TCEP, 1 x protease inhibitor cocktail, 1mM PMSF), and lysed by bead-beating (4°C, 8 cycles of 45s-on and 2min-off). The crude cell lysate was harvested by centrifugation at 950xg for 30s. 1% Triton X-100 was added at 4°C for 1.5 hr and the samples were centrifuged at 18,000xg for 10 min. 50μl was set aside as input and the remainder was incubated with anti-HA resin (Sigma, SAE0197) at 4°C for 1.5 hr. The beads were washed five times with 1 ml lysis buffer containing 0.1% Triton X-100. Bound material was eluted with 40ul of 1xSDS loading buffer.

### In vitro ubiquitination of Mga2

Yeast Uba1 was expressed in yeast and purified as previously described (*45, 46*). Yeast Rsp5 was cloned into the pK27 vector with a N-terminal His14-SUMO tag and expressed in BL21-CodonPlus cells. The cells were lysed in buffer (50 mM Tris, pH 8, 320 mM NaCl, and 10 mM imidazole supplemented with 1 mM PMSF and a protease inhibitor cocktail), and the lysate was incubated with Ni-NTA beads. Protein was eluted with elution buffer (50 mM Tris-HCl, pH 8, 150 mM NaCl, 400 mM imidazole). The His-SUMO-tag was removed with SUMO protease. The protein was concentrated, and subjected to SEC on a Superdex 200 Increase column, equilibrated in 50 mM HEPES, pH 7.4, 150 mM NaCl, 0.5 mM TCEP. Yeast Ubc4 was cloned into the pK27 vector with a His14-SUMO tag and expressed in BL21-CodonPlus cells. Ubc4 (17 kDa) was purified similarly, except that after SUMO protease digestion, the protein mixture was passed through a Ni-NTA resin to remove the SUMO-tag (14 kDa) and the SUMO protease (27 kDa). The protein was concentrated, and subjected to SEC on a Superdex 200 Increase column, equilibrated in 50 mM HEPES, pH 7.4, 150 mM NaCl, and 0.5 mM TCEP.

pRS426-plasmids encoding Flag-tagged Mga2 full-length WT, K3R (K980R, K983R, K985R) or AAKA (Rsp5 binding mutant) behind the native promoter were transformed into Δmga2 cells. Immunoprecipitation was performed with anti-Flag resin (0.5L of cell culture were used for WT and K3R samples; 1L of cell culture was used for the AAKA mutant.). Anti-Flag resin was equilibrated with ubiquitination assay buffer (50 mM HEPES, pH 7.4, 100 mM NaCl, 10% glycerol, 200 μM DTT) and divided into different tubes. Tube A contained E1 enzyme (Uba1: 200 nM), E2 enzyme (Ubc4: 1.5 μM), human ubiquitin (R&D: U-100H-10M, 60 μg), ATP/Mg2+ (5mM ATP, 10 mM MgCl2) and was incubated at RT for 10 min. Tube B contained the E3 enzyme (Rsp5: 500 nM), substrate (i.e. Mga2 bound to anti-Flag resin), ATP/Mg2+ (5mM ATP, 10 mM MgCl2). To initiate the ubiquitination reaction, the samples in tube A and B were mixed and incubated with gentle rotation at RT for 1 hr. The reaction was quenched by adding SDS loading buffer, and the samples were analyzed by SDS-PAGE followed by immunoblotting with Flag antibodies.

### CRISPR-Cas9 editing of Ole1

For tagging Ole1 at its C terminus with 2xHA in WT, Δmga2 and Δspt23 yeast cells (horizon; clone ID 4869), the guide RNA sequence (5’-3’: AGAGGTGAAATCTACGAAAC) was inserted into the pML104 plasmid by T4 ligation. The pML104 plasmid together with the donor sequence (see supplementary Table 1) was transformed into WT, Δmga2, or Δspt23 cells. Colonies on CSM-Ura plates were verified by diagnostic PCR (OLE_C-HA_F&R) and sequencing. Ole1-2xHA strains were grown in YPD medium, and total cell lysates were prepared for immunoblotting with HA and Sec62 antibodies.

### Preparation of total cell lysates

Yeast cell pellets (total OD of 8-10) were resuspended in 120 μl of lysis buffer (10 mM MOPS, 1% SDS, 8M Urea, 1x protease inhibitor cocktail, 1mM PMSF). 0.5 mm glass beads (volume∼200 μl) were added before six cycles of bead-vortexing (1min on, 1min off) at RT. 40 μl of 4xSDS loading buffer was added and mixed at 65oC for 15 min. The samples were centrifuged at 16,000xg for 2 min. The supernatants were either immediately used for SDS-PAGE or stored at −80C.

### Cycloheximide-chase experiments

Cells expressing Mga2 p90 or variants were cultured in CSM-Ura medium until an OD600 of 0.6 - 0.8. Typically, 50 ml of cell culture was harvested by centrifuging at 1,600xg for 5 min. 40ml of the supernatant was discarded, and 12 μl of 100 mg/ml cycloheximide (Sigma, 239763) was added. The cells were resuspended by brief vortexing and further cultured at 30°C. 2ml aliquots were taken every 40 min, and centrifuged at 16,000xg for 2 min. The supernatants were discarded and the pellets were frozen in liquid nitrogen and stored at −80°C. The cells were thawed on ice and total cell lysates were prepared as described above. For experiment with proteasome inhibitor, bortezomib was added to a final concentration of 80 μM, and the cells were incubated at 30°C for 2 hours before cycloheximide addition.

### Immunoblotting

Immunoprecipitated material or total cell lysates were loaded onto 4-20% Criterion TGX gels (BioRad) for SDS-PAGE. Molecular weight markers were analyzed in parallel (protein ladder: ThermoFisher, 26616). Membrane transfer was performed with the Trans-Blot Turbo system. The membrane was incubated with 5% milk/1xTBST for 2 hours at RT, before adding the primary antibody (anti-Flag, sigma, F7425; anti-HA, ChromoTek, 7c9; anti-Ubiquitin, R&D systems, MAB8595; anti-Sec62, Rapoport lab stock) overnight at 4°C. Secondary antibodies were added at room temperature for 1 hr before fluorescence scanning with the Licor system or chemiluminescence imaging with an ImageQuant800 instrument. For detecting ubiquitinated p120, HRP conjugated secondary antibody (ThermoFisher, 31460) was used. For all other blots, goat anti-rat (LICOR, #926-32219) or donkey anti-rabbit secondary antibodies (LICOR, #926-32213) were used.

### RT-qPCR

A total of 1.5 OD yeast cells were washed once with milliQ water. RNA was extracted using the YeaStar RNA Kit (Zymo research, R1002) with on-column DNase I (RNase-free, Zymo research, E1010) treatment for 15 min at RT. RNA samples were eluted with 50 μl of RNase/DNase-free water. The samples were used immediately for PCR or stored at −80C. For RT-qPCR, RNA samples were diluted to 25 ng/μl. Each reaction contained 4 μl of RNA, 0.5 μl of enzyme mix (NEB, E3005), 5 μl of one-step reaction mix, 0.08 μl of forward and reverse primers, and 0.34 μl of H2O. 5S rRNA was used as an internal control for all experiments. The data were analyzed by LightCycler 96 software. The dCq value was calculated by comparing the target gene (e.g. Ole1) to the internal control (5s rRNA), and ddCq values were converted to fold-change.

### Chromatin-immunoprecipitation (ChIP)-sequencing and qPCR

2xHA tagged p90 constructs were expressed in Δmag2 cells. Cells corresponding to a total of 70-80 OD were resuspended in 40ml of medium containing formaldehyde at a final concentration of 1%. Crosslinking was performed at RT for 15 min with constant mixing. The reaction was quenched by adding glycine to a final concentration of 0.1M. The cells were washed twice with 1xTBST, flash-frozen in liquid nitrogen, and stored at −80°C.

The cells were resuspended in ChIP buffer (50 mM HEPES, pH7.5, 140 mM Nacl, 1mM EDTA, 1% Triton x-100, 0.1% SDS, 0.1% sodium deoxycholate, 1x protease inhibitor cocktail, 1mM PMSF), lysed by bead-beating (4°C, 8 cycles of 45s-on and 2min-off). A crude cell lysate was obtained by centrifugation at 950xg for 30s, and transferred to a 1ml microtube for Covaris sonication (Covaris.520135). Each sample was sonicated for 12 min and then centrifuged at 18,000xg for 15 min. 10 μl of the supernatant was set aside as input and the remainder was incubated with anti-HA protein A magnetic beads (HA antibody: cell signaling #3724, Protein A Magnetic Beads: thermos fisher 88845) for 2 hours at 4°C. The beads were washed with ChIP buffer, buffer 500 (50 mM Tris-Hcl, pH8.0, 1mM EDTA, 1% Triton X-100, 0.1% sodium deoxycholate, 0.02% NaN3), and LiCl buffer (0.5% sodium deoxycholate, 0.02% NaN3, 1mM EDTA, 250 mM LiCl, 0.5% NP-40, 10 mM Tris-Hcl, pH8.0). The bead-bound material was eluted with 100ul of elution buffer (50 mM Tris-Hcl, pH8.0, 10 mM EDTA, 1% SDS), and incubated at 60°C overnight. 90 μl of elution buffer was added to the 10 μl input sample.

DNA was purified by ChIP DNA clean & concentrator kit (Zymo research, D5205) and stored at −80°C until library preparation or qPCR. DNA library preparation and next generation sequencing were performed by the Bauer Core Facility at Harvard. Briefly, the library was prepared using the watchmaker DNA library prep kit, checked by tapestation, and sequenced on NextSeq 1000. The data analysis was provided by the Harvard Chan Bioinformatics Core, Harvard T.H. Chan School of Public Health, Boston, MA. RRID:SCR_025373. Briefly, ChIP-seq samples were processed using the nf-core/chipseq pipeline (v2.1.0-g76e2382) executed with Nextflow (v25.10.4). Raw sequencing reads were assessed for quality using FastQC (v0.12.1). Adapter sequences and low-quality bases were trimmed using Trim Galore (v0.6.7) with Cutadapt (v3.4) with default parameters. Trimmed reads were aligned to the Saccharomyces cerevisiae reference genome (R64-1-1) using Bowtie2 (v2.5.2) with default parameters. Peaks were called using MACS3 (v3.0.1) in narrow peak mode with an effective genome size of 12,100,000 bp. Genome-wide coverage tracks were generated using bedtools genomecov (v2.30.0) with Counts Per Million (CPM) normalization. Coverage bedGraph files were sorted by genomic coordinates and converted to bigWig format using UCSC bedGraphToBigWig (v445) for visualization and downstream analysis. For qPCR using Luna® Universal qPCR Master Mix (NEB, M3003), input-DNA samples were diluted by 10-fold and 2 μl were used for each reaction; 2 μl of ChIP-DNA samples were used for each reaction. ChIP-enrichment were calculated against input-DNA samples, and then normalized to compare relative DNA binding strength.

### Ole1 promoter-URA3 reporter assay

URA3 gene under the control of Ole1 promoter sequences (whole promoter: 750bp (−750 to −1); FAR: 100bp (−576 to −477); O2R: 50bp (−356 to −307)) were cloned into the pRS413 plasmid. Empty pRS413 vector (EV) and plasmids encoding Ura3 were transformed into Δspt23 cells, and selected on CSM-HIS plates. Cells were harvested from CSM-HIS plate, serially diluted and spotted onto CSM-HIS, CSM-HIS-URA and CSM-HIS-URA + 0.05% C18:2 (Sigma L1376 + 0.2% IGEPAL-630) plates. Cells cultured in liquid CSM-HIS medium were harvested for RT-qPCR using primers targeting the Ura3 gene.

### Purification of MBP-Mga2 proteins from E. coli

The Mga2 fragment DBD-IPT-Ank (amino acids 127-842; Δ371-529) was synthesized as gene fragment (IDT) with codon optimization for bacterial expression, and cloned into the pET28 vector with an N terminal MBP-3xFlag tag. The corresponding mutants in the DBD domain were generated using Q5 mutagenesis (NEB). The proteins were expressed in BL21 codon plus competent cells (Agilent technologies, #230280). Expression was induced with 0.5 mM IPTG at 18°C for 16 hrs. The cells were lysed in lysis buffer (50 mM HEPES pH 7.4, 1mM TCEP, 500 mM NaCl, 10% glycerol, 1x protease inhibitor cocktail, and 1 mM PMSF). 1x DNase I (Sigma, 4536282001) and 10 mM MgCl2 were added to the cell suspension before lysis using an EmulsiFlex (Avestin) apparatus. The lysates were cleared by centrifugation at 44,000xg for 45 min, and the supernatant was incubated with amylose resin (NEB, E8021S) for 1 hr at 4oC. The resin was washed with 20ml of buffer 1 (50mM HEPES pH 7.4, 1mM TCEP, 1.0 M NaCl,10% glycerol), 20ml of lysis buffer (50 mM HEPES pH 7.4, 1mM TCEP, 500 mM NaCl,10% glycerol), and 20ml of buffer 2 (20 mM Tris pH7.4, 1mM TCEP, 100 mM NaCl, 10% glycerol). The MBP-fusion proteins were eluted with 20 mM Tris pH7.4, 1mM TCEP, 100 mM NaCl, 10% glycerol, 30 mM maltose. The eluates were concentrated and loaded onto a Heparin column (Cytiva, 17040601) equilibrated with buffer 2, and then eluted over 1 hr with a linear salt gradient using buffer 3 (20mM Tris pH7.4, 1mM TCEP, 2M NaCl, 10% glycerol). Peak fractions were collected for SDS-PAGE, concentrated, and stored at −80°C. The Mga2 fragment containing only the DBD was cloned into the pET28 vector with an N terminal His-MBP tag, and was expressed and purified in a similar manner.

### DNA gel shift assays

Cy3-O2R oligos (forward and reverse, IDT) were diluted to 1.25 μM using buffer A (20mM HEPES, 100 mM Nacl, pH7.4, 10% glycerol, 1mM TCEP). Equal volumes of forward and reverse oligos were mixed and incubated at 95°C for 5 min, before cooling to RT over 30 min. The Mga2 fragments were added at a final concentration of 0.5 μM to the Cy3-O2R sample (final concentration of dsDNA 0.1 μM) in binding buffer (buffer A + 5 mM MgCl2). The mixture was then incubated at RT for 30 min before electrophoresis in a 5% native polyacrylamide gel at 100V for 60 min in 0.5xTBE buffer (ice-bath). The gel was directly scanned on a Licor imager to detect the Cy3 signal. Non-fluorescent competitors were added 15 min before Cy3-O2R to a final concentration of at 0.4 μM. Gel shift assays with MBP-DBD alone were performed similarly.

### Fluorescence microscopy

Cells expressing GFP-Mga2 (p90 or p120) and NAB2-mCherry were cultured in CSM-Ura medium to OD600 of 0.6-0.8. Cells stably expressing NAB2-mCherry were generated by deleting the TRP1 gene from BY4741 cells using pFA6a-natMX, and introducing the gene coding for NAB2-mCherry using Trp1 as a selection marker (Addgene: #133648). The cells were resuspended to a density of 8-10 OD/ml, and 10 μl were applied to a gelatin pad on an imaging glass slide. A cover slip was added and the sample sealed with Valap solution (Vaseline: lanolin: paraffin wax=1:1:1). Z-stack images were acquired on an Yokagawa CSU-X1 spinning disk confocal microscope (CITE, Harvard Medical School) with 20% laser output and an exposure time of 400 ms at 488 nm wavelength for GFP and 561 nm for mCherry. For lipid droplet imaging, cells from an overnight culture were harvested and resuspended to a density of 8-10 OD/ml, and stained with 5 μM BODIPY 493/503 (MCE, HY-W090090) for 30 min at 30oC. The cells were washed once with fresh medium and then prepared for imaging with 20% laser output, an exposure time of 400 ms, and a wavelength of 488 nm. Fluorescence images were analyzed with ImageJ software.

### Flow cytometry measurement of Lipid droplets

Mga2 p90 fragments (residue 1-626) were expressed in Δmga2 cells from the pRS416 plasmid under a TEF1 promoter. Cells from overnight cultures were incubated at 3OD/ml with 5 μM of BODIPY 493/503 for 30 min at 30°C under shaking. The cells were washed once with fresh CSM-Ura medium, and diluted to 0.2 OD/ml. They were analyzed with an Attune Flow Cytometer with 488nm excitation and BL1 emission filters (530/30). A total of 10,000 single cells/events were collected and quantified for each sample after gating. Unstained cells were used as a control to account for autofluorescence. Flow cytometry data were analyzed and visualized in FlowJo v10.7.1.

### GC-MS analysis of cellular acyl chain composition

80 OD of cells were harvested from an overnight cell culture and washed once with H2O. The cells were lysed in 20mM HEPES, 100 mM Nacl, pH7.4, 10% glycerol, 1mM TCEP by bead-beating. Lipid extraction from cell lysates (200 μl) were performed using the Bligh and Dyer extraction protocol (chloroform: methanol=1:2) (*47*). The chloroform phase was collected and dried using a stream of nitrogen gas. 1 ml of 1 M methanolic HCl (Sigma, 90964) containing 5% v/v 2,2-dimethoxypropane (Sigma, D136808) was added together with 200 μg of d35-C18:0 internal standard (Cayman, 9003318) to prepare fatty acid methyl esters. The samples were incubated at 80°C for 1hr and cooled to room temperature. 2ml of 0.9% NaCl and 1ml of hexane were added and vortexed briefly. The samples were centrifuged at 1000 x g for 2 min, and the upper hexane phase was collected. The GC-MS analysis (signature spectrum m/z=74) was performed by the Harvard Center for Mass Spectrometry. The relative abundance of C18:0 and C18:1 was quantified using the internal standard.

### Yeast viability assays

The mag2spt23 double knock-out (DKO) (*48*) strain was a gift from Dr. Robert Ernst (Saarland University, Germany). The DKO strain was maintained and cultured in YPD medium + 1% (w/v) IGEPAL-630 (Sigma, I8896) + 0.05% (w/v) sodium oleate (Sigma, O3880). Plasmid transformation was performed with the LiAC/PEG3350 method (*49*). The cells were grown at 30oC for three days on CSM-Ura plates containing 0.2% IGEPAL-630 and 0.05% (w/v) sodium oleate. To check viability in the absence of oleate, cells were harvested from oleate-containing medium, suspended in sterile H2O, and diluted to an OD600 of 1.0 before serial dilution. The cells were spotted or spread onto CSM-Ura plates lacking or containing oleate, and incubated at 30°C for three days.

A yeast strain lacking lipid droplets (4KO-LD: are1Δ are2Δ lro1Δ dga1Δ) was a gift from Dr. Amit Joshi (University of Tennessee, Knoxville, USA) and was previously described by Choudhary et al (*50*). Mga2 was deleted from the 4KO-LD cells using a hygromycin-resistance cassette (addgene: 228258). The deletion was verified by diagnostic PCR. An empty vector (pRS416-TEF1 promoter) or plasmids (pRS416-TEF1 promoter) encoding wild-type p90 (residue 1-626) or the mutants KR or 3A were transformed into Δmga2 (+Lipid droplets in Fig.6F) and 4KO-LD/Δmga2 (-Lipid droplets in Fig.6F) strains. Transformants were incubated on CSM-URA plates at 30°C for 3 days.

### Statistical Analysis

Statistical analysis among groups was performed using Prism (GraphPad Software, RRID: SCR_002798; http://www.graphpad.com) with One-way ANOVA test. Results were indicated in the following manner: * for P < 0.05, ** for P < 0.01, *** for P < 0.001, and **** for P < 0.0001, where P < 0.05 is considered as significantly different.

## Acknowledgments

We thank Hao Li for gifts of purified Uba1, Rsp5, and Ubc4 proteins; Shannan Ho Sui from the Harvard Chan Bioinformatics Core for help with ChIP-seq data analysis; Rudi Pisa, Amit Joshi, and Robert Ernst for sharing yeast reagents used in this study; and Shashank Rao for assistance with Ole1 strain generation. We thank the Harvard Cell Biology Education and Fellowship Fund (Goldberg Fellowship awarded to N.W.) for the funding support.

## Author contributions

Conceptualization: N.W., T.A.R.

Methodology: N.W., J.K., J.A.P., C.N.

Investigation: N.W., J.K., J.A.P., C.N.

Visualization: N.W., J.K., J.A.P., C.N.

Supervision: T.A.R., S.P.G.

Writing—original draft: T.A.R.

Writing—review & editing: N.W., T.A.R.

## Data, code, and materials availability

All data and code needed to evaluate and reproduce the results in the paper are present in the paper and/or the Supplementary Materials. ChIP-seq data will be deposited at GEO. Mass spectrometry data will be deposited to the ProteomeXchange Consortium. All materials (plasmids and strains) will be made available upon request to T.A.R.

## REFERENCES

1. I. Shimomura, H. Shimano, B. S. Korn, Y. Bashmakov, J. D. Horton, Nuclear sterol regulatory element-binding proteins activate genes responsible for the entire program of unsaturated fatty acid biosynthesis in transgenic mouse liver. J Biol Chem 273, 35299–35306 (1998).

2. J. E. Stukey, V. M. McDonough, C. E. Martin, The OLE1 gene of Saccharomyces cerevisiae encodes the delta 9 fatty acid desaturase and can be functionally replaced by the rat stearoyl-CoA desaturase gene. J Biol Chem 265, 20144–20149 (1990).

3. J. Y. Choi, J. Stukey, S. Y. Hwang, C. E. Martin, Regulatory elements that control transcription activation and unsaturated fatty acid-mediated repression of the Saccharomyces cerevisiae OLE1 gene. J Biol Chem 271, 3581–3589 (1996).

4. Y. Jiang, M. J. Vasconcelles, S. Wretzel, A. Light, L. Gilooly, K. McDaid, C. S. Oh, C. E. Martin, M. A. Goldberg, Mga2p processing by hypoxia and unsaturated fatty acids in Saccharomyces cerevisiae: impact on LORE-dependent gene expression. Eukaryot Cell 1, 481–490 (2002).

5. K. E. Kwast, P. V. Burke, B. T. Staahl, R. O. Poyton, Oxygen sensing in yeast: evidence for the involvement of the respiratory chain in regulating the transcription of a subset of hypoxic genes. Proc Natl Acad Sci U S A 96, 5446–5451 (1999).

6. S. Zhang, T. J. Burkett, I. Yamashita, D. J. Garfinkel, Genetic redundancy between SPT23 and MGA2: regulators of Ty-induced mutations and Ty1 transcription in Saccharomyces cerevisiae. Mol Cell Biol 17, 4718–4729 (1997).

7. S. Zhang, Y. Skalsky, D. J. Garfinkel, MGA2 or SPT23 is required for transcription of the delta9 fatty acid desaturase gene, OLE1, and nuclear membrane integrity in Saccharomyces cerevisiae. Genetics 151, 473–483 (1999).

8. T. Hoppe, K. Matuschewski, M. Rape, S. Schlenker, H. D. Ulrich, S. Jentsch, Activation of a membrane-bound transcription factor by regulated ubiquitin/proteasome-dependent processing. Cell 102, 577–586 (2000).

9. M. Rape, T. Hoppe, I. Gorr, M. Kalocay, H. Richly, S. Jentsch, Mobilization of processed, membrane-tethered SPT23 transcription factor by CDC48(UFD1/NPL4), a ubiquitin-selective chaperone. Cell 107, 667–677 (2001).

10. S. Ballweg, R. Ernst, Control of membrane fluidity: the OLE pathway in focus. Biol Chem 398, 215–228 (2017).

11. J. Ye, R. A. DeBose-Boyd, Regulation of cholesterol and fatty acid synthesis. Cold Spring Harb Perspect Biol 3, (2011).

12. R. Covino, S. Ballweg, C. Stordeur, J. B. Michaelis, K. Puth, F. Wernig, A. Bahrami, A. M. Ernst, G. Hummer, R. Ernst, A Eukaryotic Sensor for Membrane Lipid Saturation. Mol Cell 63, 49–59 (2016).

13. N. Shcherbik, T. Zoladek, J. T. Nickels, D. S. Haines, Rsp5p is required for ER bound Mga2p120 polyubiquitination and release of the processed/tethered transactivator Mga2p90. Curr Biol 13, 1227–1233 (2003).

14. N. Shcherbik, Y. Kee, N. Lyon, J. M. Huibregtse, D. S. Haines, A single PXY motif located within the carboxyl terminus of Spt23p and Mga2p mediates a physical and functional interaction with ubiquitin ligase Rsp5p. J Biol Chem 279, 53892–53898 (2004).

15. Y. Saeki, T. Kudo, T. Sone, Y. Kikuchi, H. Yokosawa, A. Toh-e, K. Tanaka, Lysine 63-linked polyubiquitin chain may serve as a targeting signal for the 26S proteasome. EMBO J 28, 359–371 (2009).

16. W. Piwko, S. Jentsch, Proteasome-mediated protein processing by bidirectional degradation initiated from an internal site. Nat Struct Mol Biol 13, 691–697 (2006).

17. L. Lin, S. Ghosh, A glycine-rich region in NF-kappaB p105 functions as a processing signal for the generation of the p50 subunit. Mol Cell Biol 16, 2248–2254 (1996).

18. D. A. Kraut, Slippery substrates impair ATP-dependent protease function by slowing unfolding. J Biol Chem 288, 34729–34735 (2013).

19. Y. Jiang, M. J. Vasconcelles, S. Wretzel, A. Light, C. E. Martin, M. A. Goldberg, MGA2 is involved in the low-oxygen response element-dependent hypoxic induction of genes in Saccharomyces cerevisiae. Mol Cell Biol 21, 6161–6169 (2001).

20. Y. Nakagawa, S. Sugioka, Y. Kaneko, S. Harashima, O2R, a novel regulatory element mediating Rox1p-independent O(2) and unsaturated fatty acid repression of OLE1 in Saccharomyces cerevisiae. J Bacteriol 183, 745–751 (2001).

21. M. J. Vasconcelles, Y. Jiang, K. McDaid, L. Gilooly, S. Wretzel, D. L. Porter, C. E. Martin, M. A. Goldberg, Identification and characterization of a low oxygen response element involved in the hypoxic induction of a family of Saccharomyces cerevisiae genes. Implications for the conservation of oxygen sensing in eukaryotes. J Biol Chem 276, 14374–14384 (2001).

22. H. Richly, M. Rape, S. Braun, S. Rumpf, C. Hoege, S. Jentsch, A series of ubiquitin binding factors connects CDC48/p97 to substrate multiubiquitylation and proteasomal targeting. Cell 120, 73–84 (2005).

23. B. Ho, A. Baryshnikova, G. W. Brown, Unification of Protein Abundance Datasets Yields a Quantitative Saccharomyces cerevisiae Proteome. Cell Syst 6, 192–205 e193 (2018).

24. G. Ghosh, G. van Duyne, S. Ghosh, P. B. Sigler, Structure of NF-kappa B p50 homodimer bound to a kappa B site. Nature 373, 303–310 (1995).

25. C. W. Muller, F. A. Rey, M. Sodeoka, G. L. Verdine, S. C. Harrison, Structure of the NF-kappa B p50 homodimer bound to DNA. Nature 373, 311–317 (1995).

26. J. A. Fleming, E. S. Lightcap, S. Sadis, V. Thoroddsen, C. E. Bulawa, R. K. Blackman, Complementary whole-genome technologies reveal the cellular response to proteasome inhibition by PS-341. Proc Natl Acad Sci U S A 99, 1461–1466 (2002).

27. S. Bhattacharya, N. Shcherbik, J. Vasilescu, J. C. Smith, D. Figeys, D. S. Haines, Identification of lysines within membrane-anchored Mga2p120 that are targets of Rsp5p ubiquitination and mediate mobilization of tethered Mga2p90. J Mol Biol 385, 718–725 (2009).

28. L. Holm, A. Laiho, P. Toronen, M. Salgado, DALI shines a light on remote homologs: One hundred discoveries. Protein Sci 32, e4519 (2023).

29. M. van Kempen, S. S. Kim, C. Tumescheit, M. Mirdita, J. Lee, C. L. M. Gilchrist, J. Soding, M. Steinegger, Fast and accurate protein structure search with Foldseek. Nat Biotechnol 42, 243–246 (2024).

30. J. Jumper, R. Evans, A. Pritzel, T. Green, M. Figurnov, O. Ronneberger, K. Tunyasuvunakool, R. Bates, A. Zidek, A. Potapenko, A. Bridgland, C. Meyer, S. A. A. Kohl, A. J. Ballard, A. Cowie, B. Romera-Paredes, S. Nikolov, R. Jain, J. Adler, T. Back, S. Petersen, D. Reiman, E. Clancy, M. Zielinski, M. Steinegger, M. Pacholska, T. Berghammer, S. Bodenstein, D. Silver, O. Vinyals, A. W. Senior, K. Kavukcuoglu, P. Kohli, D. Hassabis, Highly accurate protein structure prediction with AlphaFold. Nature 596, 583–589 (2021).

31. K. H. Vousden, C. Prives, Blinded by the Light: The Growing Complexity of p53. Cell 137, 413–431 (2009).

32. B. Yariv, E. Yariv, A. Kessel, G. Masrati, A. B. Chorin, E. Martz, I. Mayrose, T. Pupko, N. Ben-Tal, Using evolutionary data to make sense of macromolecules with a “face-lifted” ConSurf. Protein Sci 32, e4582 (2023).

33. W. Wen, J. L. Meinkoth, R. Y. Tsien, S. S. Taylor, Identification of a signal for rapid export of proteins from the nucleus. Cell 82, 463–473 (1995).

34. N. Raj, L. D. Attardi, The Transactivation Domains of the p53 Protein. Cold Spring Harb Perspect Med 7, (2017).

35. P. H. Kussie, S. Gorina, V. Marechal, B. Elenbaas, J. Moreau, A. J. Levine, N. P. Pavletich, Structure of the MDM2 oncoprotein bound to the p53 tumor suppressor transactivation domain. Science 274, 948–953 (1996).

36. L. Sandager, M. H. Gustavsson, U. Stahl, A. Dahlqvist, E. Wiberg, A. Banas, M. Lenman, H. Ronne, S. Stymne, Storage lipid synthesis is non-essential in yeast. J Biol Chem 277, 6478–6482 (2002).

37. J. A. Johnston, E. S. Johnson, P. R. Waller, A. Varshavsky, Methotrexate inhibits proteolysis of dihydrofolate reductase by the N-end rule pathway. J Biol Chem 270, 8172–8178 (1995).

38. A. Peth, J. A. Nathan, A. L. Goldberg, The ATP costs and time required to degrade ubiquitinated proteins by the 26 S proteasome. J Biol Chem 288, 29215–29222 (2013).

39. G. A. Collins, A. L. Goldberg, The Logic of the 26S Proteasome. Cell 169, 792–806 (2017).

40. P. Rice, I. Longden, A. Bleasby, EMBOSS: the European Molecular Biology Open Software Suite. Trends Genet 16, 276–277 (2000).

41. C. E. Martin, C. S. Oh, Y. Jiang, Regulation of long chain unsaturated fatty acid synthesis in yeast. Biochim Biophys Acta 1771, 271–285 (2007).

42. M. Muratani, W. P. Tansey, How the ubiquitin-proteasome system controls transcription. Nat Rev Mol Cell Biol 4, 192–201 (2003).

43. M. C. J. Yip, N. O. Bodnar, T. A. Rapoport, Ddi1 is a ubiquitin-dependent protease. Proc Natl Acad Sci U S A 117, 7776–7781 (2020).

44. J. A. Paulo, Sample preparation for proteomic analysis using a GeLC-MS/MS strategy. J Biol Methods 3, (2016).

45. Z. Ji, H. Li, D. Peterle, J. A. Paulo, S. B. Ficarro, T. E. Wales, J. A. Marto, S. P. Gygi, J. R. Engen, T. A. Rapoport, Translocation of polyubiquitinated protein substrates by the hexameric Cdc48 ATPase. Mol Cell 82, 570–584 e578 (2022).

46. H. Li, Z. Ji, J. A. Paulo, S. P. Gygi, T. A. Rapoport, Bidirectional substrate shuttling between the 26S proteasome and the Cdc48 ATPase promotes protein degradation. Mol Cell 84, 1290–1303 e1297 (2024).

47. R. Schneiter, G. Daum, Extraction of yeast lipids. Methods Mol Biol 313, 41–45 (2006).

48. S. Ballweg, E. Sezgin, M. Doktorova, R. Covino, J. Reinhard, D. Wunnicke, I. Hanelt, I. Levental, G. Hummer, R. Ernst, Regulation of lipid saturation without sensing membrane fluidity. Nat Commun 11, 756 (2020).

49. R. D. Gietz, R. A. Woods, Yeast transformation by the LiAc/SS Carrier DNA/PEG method. Methods Mol Biol 313, 107–120 (2006).

50. V. Choudhary, O. El Atab, G. Mizzon, W. A. Prinz, R. Schneiter, Seipin and Nem1 establish discrete ER subdomains to initiate yeast lipid droplet biogenesis. J Cell Biol 219, (2020).

