## Supplementary Files for "Activation and inactivation pathways of a p53-like transcription factor govern lipid homeostasis in yeast"

Neng Wan *et al.*

**This PDF file includes:**

Figs. S1 to S4
Tables S1 to S3

**Figure S1**

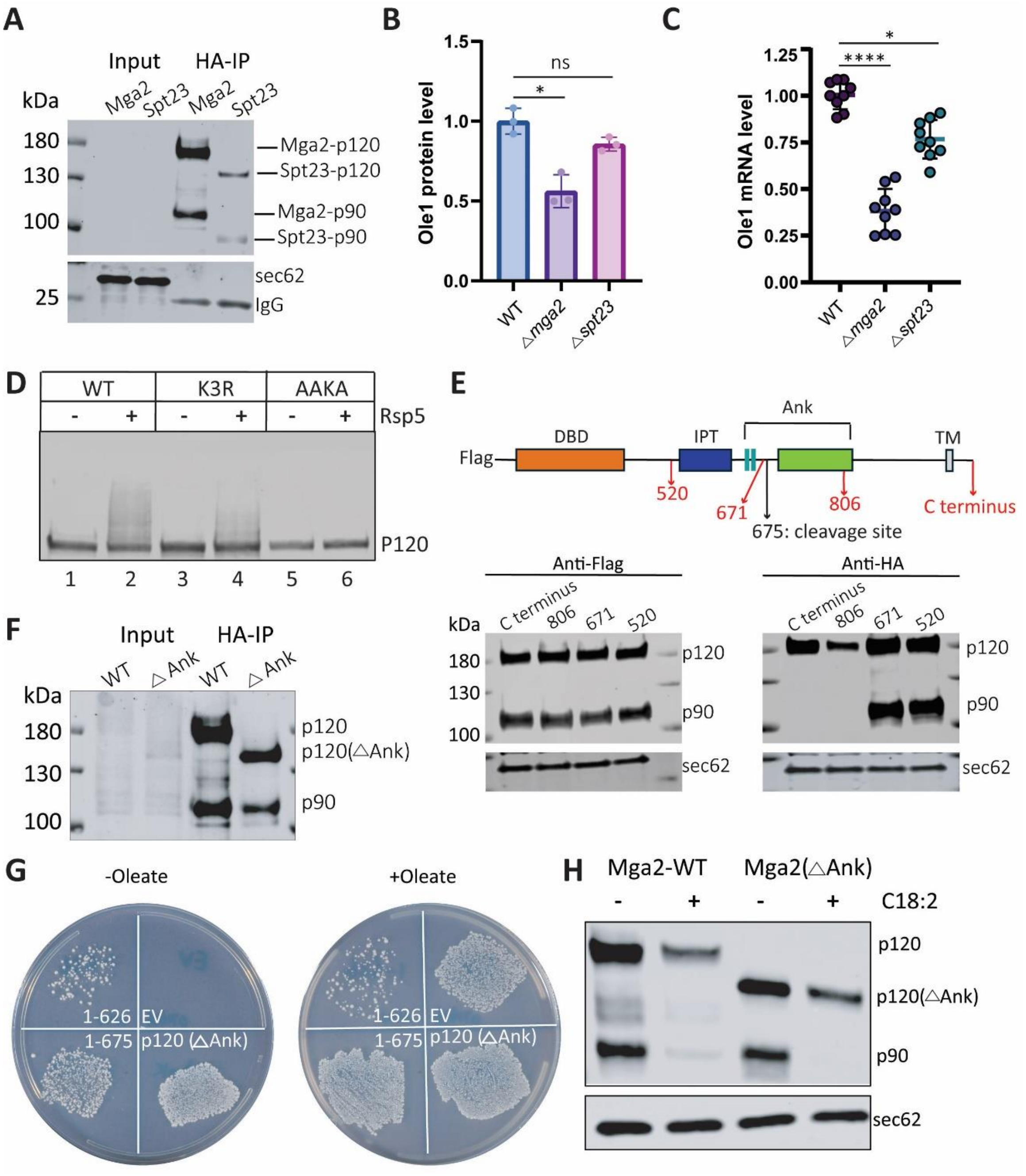


**Figure S1. Processing of the Mga2 precursor.**

(**A**) Mga2 or Spt23 with N-terminal 2xHA tags were expressed from the chromosome. Triton-solubilized cell lysates were incubated with HA-antibody beads and the bound material was analyzed by SDS-PAGE and immunoblotting with HA antibodies. Sec62 and immunoglobulin (IgG) served as loading controls. (**B**) RNA was extracted from wild-type (WT) cells or cells lacking Mga2 or Spt23 (Δmga2 or Δspt23) and the level of Ole1 mRNA determined by RT-qPCR. The levels of Ole1 transcripts are expressed relative to those of WT. (**C**) Ole1 was tagged at the C-terminus with 2xHA epitopes and expressed in WT, ∆mga2, or ∆spt23 cells. Cell lysates were analyzed by SDS-PAGE and immunoblotting with HA antibodies. The intensity of the Ole1-2xHA bands was quantified and is expressed relative to that of WT. (**D**) Full-length wild-type (WT) Mga2 or mutants with three lysine residues mutated (K3R: K980R, K983R, K985R) or defective in Rsp5 binding (AAKA) were Flag-tagged at the N-terminus and over-expressed in ∆mga2 cells. Cell lysates were subjected to immunoprecipitation with anti-Flag beads. After washing, the beads were incubated with purified Rsp5 and the bead-bound material analyzed by SDS-PAGE and immunoblotting with Flag antibodies. Controls were performed in the absence of Rsp5. (**E**) The HA epitope was inserted at the indicated positions into Flag-tagged full-length Mga2, and the proteins were over-expressed in ∆mga2 cells. Cell lysates were analyzed by immunoblotting with Flag and HA antibodies. Blotting for Sec62 served as a loading control. (**F**) Full-length Mga2 carrying N-terminal 2xHA tags was expressed from the chromosome. Cell lysates were subjected to immunoprecipitation with HA-antibody beads. The samples were analyzed by SDS-PAGE and immunoblotting with HA antibodies. (**G**) Mga2 fragments containing residues 1-675 or 1-626 with an N-terminal Flag-tag were over-expressed in cells lacking both Mga2 and Spt23. Where indicated, cells expressed from a centromeric plasmid a p120 variant that lacks the Ank domain. An empty vector (EV) was used as control. The cells were grown on plates lacking or containing oleate at 30°C for three days. (**H**) As in (F) but with addition of unsaturated fatty acid C18:2, as indicated.

**Figure S2**

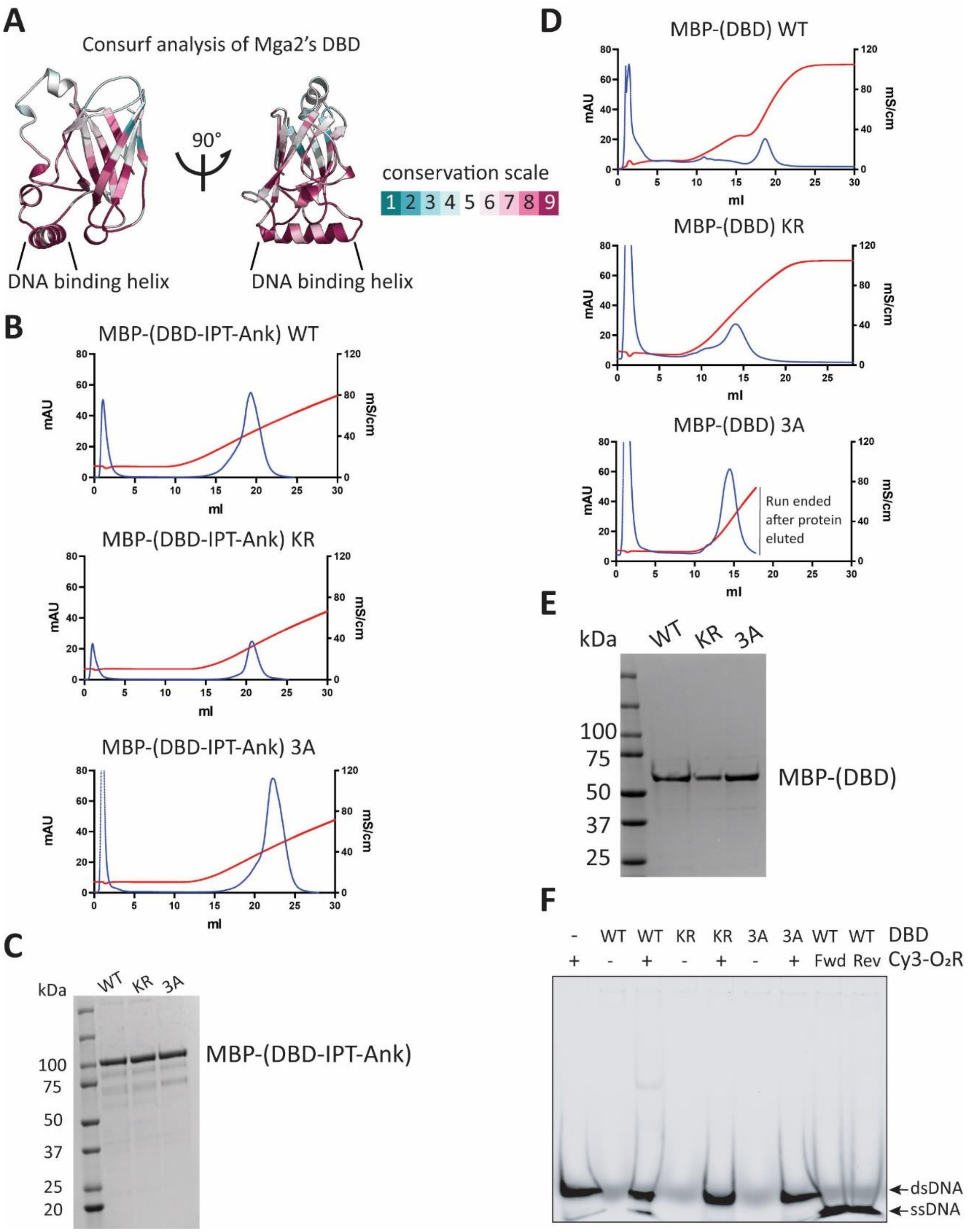


**Figure S2. Structural conservation of Mga2's DBD and purification of DBD-containing proteins.**

(**A**) Conservation of the DBD as analyzed by the program Consurf. The AlphaFold-predicted structure was used as input. The scale of conservation is shown on the right. Note that the

DNA binding helix is highly conserved. (**B**) Mga2 fragments spanning the region from the DBD to the Ank domain were expressed as fusions with the maltose-binding protein (MBP) in E. coli (MBP-(DBD-IPT-Ank). The DBD domain was either wild-type (WT) or contained the KR or 3A mutations. The proteins were purified on an amylose resin and applied to a heparin column. Bound protein was eluted with a linear salt gradient. (**C**) The purified proteins in b were subjected to SDS-PAGE and Coomassie-blue staining. (**D**) As in (B), but with MBP-fusions of the DBD alone. (**E**) As in (C), but with MBP-DBD fusions. (**F**) Gel-shift assays using MBP-DBD fusions and Cy3-labeled O2R DNA. The experiment was performed as in Fig. 2D.

**Figure S3**

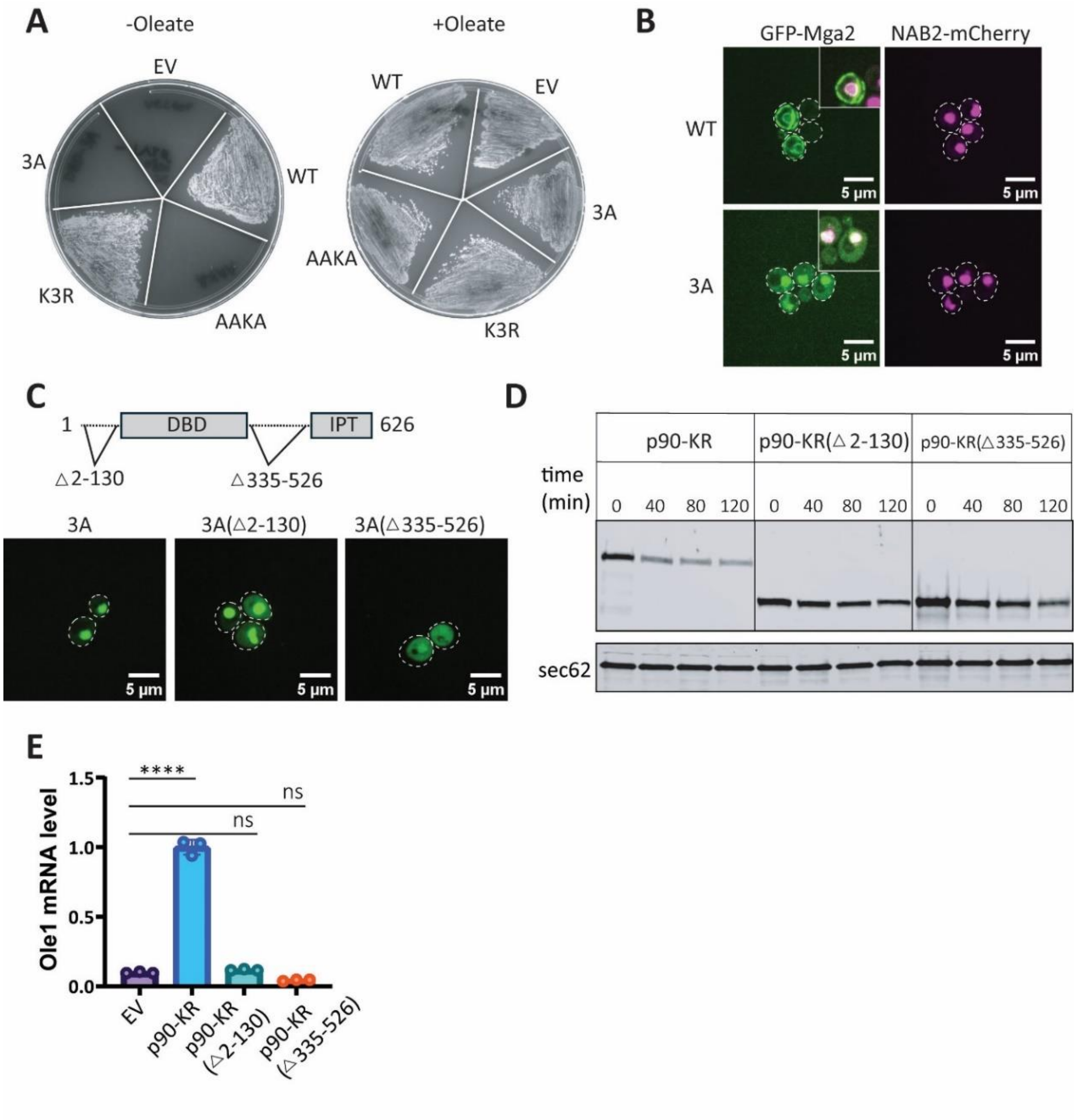


**Figure S3. Degradation of p90 requires nuclear import and DNA binding.**
(**A**) Full-length wild-type (WT) Mga2 or the indicated mutants were expressed from the native promoter on a centromeric plasmid in cells lacking Mga2 and Spt23. The cells were grown on plates lacking or containing oleate at 30°C for three days. Controls cells received an empty vector (EV). (**B**) GFP-tagged full-length Mga2 or the corresponding 3A mutant were expressed from the native promoter on a 2m plasmid. The cells also stably expressed NAB2-mCherry as a nuclear marker. GFP and mCherry fluorescence was analyzed by confocal microscopy. The inset shows merged images. The boundaries of the cells are indicated. (**C**) GFP-tagged p90 (residues 1-626) containing the 3A mutation (3A) or variants that lack residues 2-130 or 335-526 (3A (Δ2-130) and 3A (Δ335-526), respectively) were expressed from the native promoter on a 2μ plasmid in Δmga2 cells. GFP fluorescence was analyzed by confocal microscopy. The boundaries of the cells are indicated. (**D**) HA-tagged p90 carrying the KR mutation (p90-KR; residue 1-626) or variants lacking residues 2-130 or 335-526 (p90-KR (Δ2-130) and p90-KR

(∆335-526) were over-expressed in ∆mga2 cells. Cycloheximide was added, and samples taken at different time points were analyzed by SDS-PAGE and immunoblotting with HA antibodies. (**E**) RNA was extracted from the indicated cells and Ole1 mRNA levels were quantified by RT-qPCR. The levels of Ole1 transcripts are expressed relative to those of p90-KR.

**Figure S4**

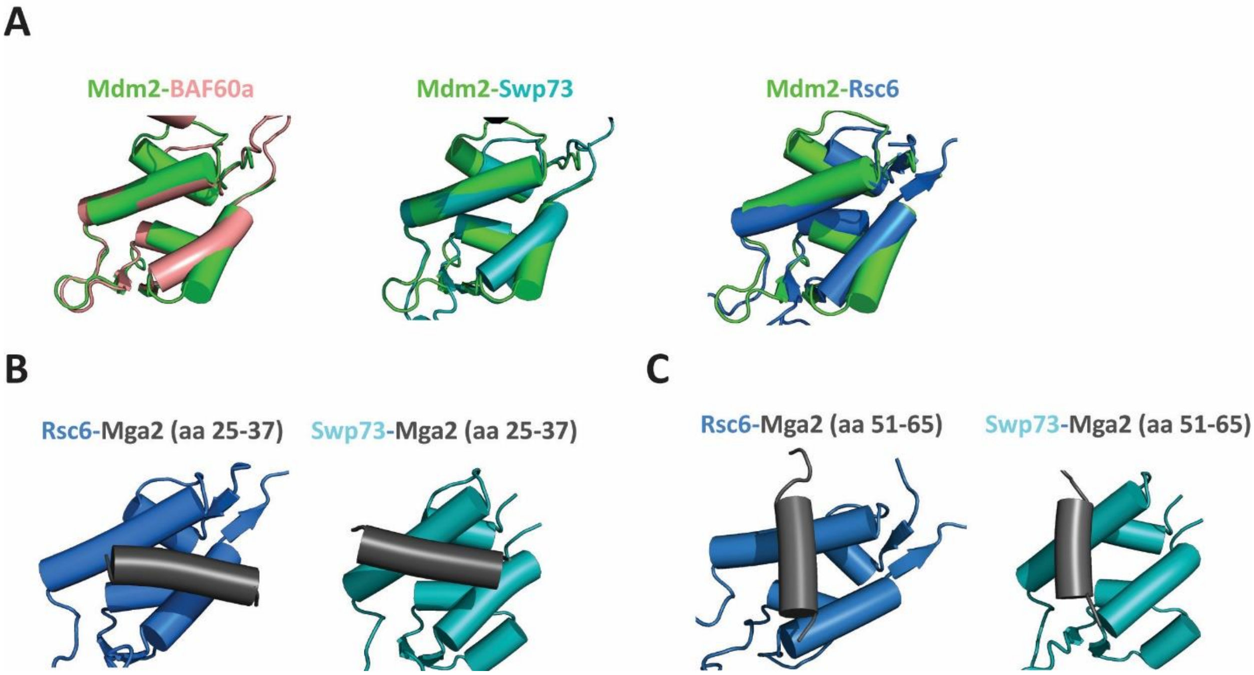


**Figure S4. Predicted interactions of Mga2 with chromatin remodeling components.**
(**A**) Structural comparison of the TAD1-binding domain of Mdm2 with domains of chromatin remodeling complexes. BAF60a (also called SMARCD1) and Swp73 are components of the human and yeast SWI/SNF complexes, respectively (r.m.s.d: 0.91Å and 0.77Å), and Rsc6 is a component of the yeast RSC complex (r.m.s.d: 2.11Å). (**B**) AlphaFold predictions of complexes of the TAD-like sequence of Mga2 (dark grey, residues 25-37) and the binding domains of Rsc6 and Swp73 (iPTM=0.53 and 0.62, respectively). (**C**) AlphaFold predictions of complexes of residues 51-65 of Mga2 (dark grey) and the binding domains of Rsc6 and Swp73 (iPTM=0.64 and 0.59, respectively).

### Table S1: primers used in the study

| Name | Sequence (5'-3') |
|---|---|
| GU_Mga2_F | AGGAGAAAAAACCCGGGTACCATGGACTACAAAGACCATG |
| GU_Mga2_R | CCGCTTATTTAGAAGTGTCGAATTCCTAACTGACAATTAAATCGTTC |
| GL_RSP5_F | AGTAAGAATTTTTGAAAGGGGATCCATGTACCCATACGATGTTCCAGATTACGCTCCTTCATCCATATCCGTC |
| GL_RSP5_R | AAAGCTTGCATGCCTGCAGGTCGACTCATTCTTGACCAAACCCTATGGTTTCTTCC |
| pET28-DBD_F | CCTGTATTTCCAGGGAGAAGGATCCTACATAAATCCTTTAGATTACCTTAAAG |
| pET28-DBD_R | TGGTGGTGGTGCTCGAGTGCGGCCGCTCATGTCGTAAACTTACCCAACAG |
| K254R_F | CGCTAGAAGGCGCTCAGGTATT |
| K254R_R | GCTCTTCTTTGTTCTCGC |
| 3A_F | TGCAGCTGCGTCAGGTATTGCAGACAATTTACTCTGGT |
| 3A_R | GCGGCTCTTCTTTGTTCTCGCTTAAC |
| pET28_MBP_F | ATTTCCAGGGAGAAGGATCCGACTACAAAGACCATGACGG |
| MBP_R | CCGTCATGGTCTTTGTAGTCGGATCCTTCTCCCTGGAAAT |
| pET28a_Mga2_F | CACCAAATTTACAGACAGCGTATAAGCTCGAGCACCACCA |
| Mga2_R | CGCTGTCTGTAAATTTGGTGCCGTTCTGGAATTCTTCAAA |
| MBP-K254R_F | CGCGTCGCCGATCAGGTATC |
| MBP-K254R_R | CAGCACGACGCTGCTC |
| MBP-3A_F | CGGCTTCAGGTATCGCAGATAACTTGC |
| MBP-3A_R | CTGCCGCAGCACGACGCTG |
| Cas9_donor_2xHA-Mga2 | TTATTGAAGGTCATTTTGGCGAACAGAACATTTCGTTATGTACCCTTACGATGTGCCCGATTACGCCTACCCTTACGATGT GCCCGATTACGCC CAGCAGAACAGTGAGTTCTTAACT GAA ACACCTGGAAGCGACCCTCATATATCTCAATTGCACGCGA |
| Cas9_donor_2xHA-Spt23 | AACGACTAATCACAACAGTAGTACACCACTGAAAATGATGTACCCTTACGATGTGCCCGATTACGCCTACCCTTACGATGT GCCCGATTACGCC AGT GGC ACA GCA AACGTTTCCTCGATGCTCCACAGCTATAGCGCCAACATAC |
| Cas9_donor_2xHA-Mga2 (ΔAnk) | ATTGAGATCAATGACAATAAAAAGGCCATATTTACCTATGTTGATAAGTTTACCGATAGTGTAGAAACAGACAGTAATTAT TCTATTAGC |
| Cas9_donor_Ole1-2xHA | CTGTTATCAAGGAAAGTAAGAACTCTGCTATTAGAATGGCTAGTAAGAGAGGTGAAATCTACGAAACTGCTAAGTTCTTTT ATCCATACGACGTACCAGATTATGCTTACCCTTACGATGTGCCCGATTACGCCTAAGTATCACATTACAATAACAAAACTG CAACTACCATAAAAAAAAATTGAAAAATCAT |
| Mga2_check_F | CTGGCACTTTGTCTCAGGGTTAGTG |
| Mga2_check_R | TTTACATTTAAAGTATATACATTATCGTTACGCTGAC |

| | |
|---|---|
| Spt23_check_F | TTTCGATGGTGTACGTTGTATCAG |
| Spt23_check_R | AAAGAAGATAAAATTCAAAAATGCAAAATAATAAAAAATGAATC |
| OLE_ C-HA_F | GATCTTCTCCGCTCACTGGCCATTGA |
| OLE_ C-HA_R | AGCCGCCTTGCATGGTGCTTTGTCATT |
| Ole1P_750_F1 | TGGAGCTCCACCGCGGTGGCGGCCGCAAGGATTAGCGGATATGTAGTTCC |
| Ole1P_750_R1 | CTTTCGACATCTTTGTTGTAATGTTTTAGTGCTG |
| Ole1P_750_F2 | TACAACAAAGATGTCGAAAGCTACATATAAGGAACG |
| Ole1P_750_R2 | GTGGCGCGCCTTAGTTTTGCTGGCCGCATC |
| Ole1p_FAR_F1 | TGGAGCTCCACCGCGGTGGCGGCCGCGGGCATGTCCCGGGGTTA |
| Ole1p_FAR_R1 | CTTTCGACATGCTGGGATACCCGAAATAGCTC |
| Ole1p_FAR_F2 | GTATCCCAGCATGTCGAAAGCTACATATAAGGAAC |
| Ole1p_O2R_F1 | TGGAGCTCCACCGCGGTGGCGGCCGCTTCTTTCGGACGTTGAACACTC |
| Ole1p_O2R_R1 | CTTTCGACATTAGAAGCACACCTGGTTGGG |
| Ole1p_O2R_F2 | TGTGCTTCTAATGTCGAAAGCTACATATAAGGAAC |
| 2xHA_Mga2_F | TATCCATACGACGTACCAGATTATGCTCAGCAGAACAGTGAGTTC |
| Mga2_426_R | CGAATTCCTGCAGCCCGGGGGATCCTTTACATTTAAAGTATATACATTATCGTTAC |
| F131_F | TTTGGAAGAAAGGAGACG |
| 2xHA_Q2_R | AGCATAATCTGGTACGTCG |
| TAD-4E_F | TGAGGAGCTGAACGGGTCTCCCATG |
| TAD-4E_R | TCGTCTTCTTCCTGTGATTCCATTACGCTATTCG |
| Mga2-416_F | TGGAGCTCCACCGCGGTGGCGGCCGCATCTTTGCTTCAAAGATTTTTTG |
| Mga2-416_R | CGAATTCCTGCAGCCCGGGGGATCCTTTACATTTAAAGTATATACATTATCGTTAC |
| AAKA_F | AAAGCTGAGGATCTGTTCCCGTT |
| AAKA_R | TGCTGCGTCATCATTAATTCGATGTAAAACTCT |
| K985_F | CGATTGCGAACCACAAATCAAGACAGTATTGTG |
| K980_R | ATCATCTCGACCCCAAGACAACGG |
| Ank_F | AAGTTTACCGATAGTGTAG |
| Ank_R | AACATAGGTAAATATGGCC |
| Mga2-TEF1_F | TAAGTTTTCTAGAACTAGTGGATCCATGGATTATAAAGATGATGATGATAAAC |
| Mga2-TEF1_R | GACATAACTAATTACATGACTCGAGCTAACTGACAATTAAATCGTTCAAC |
| TEF1_626_R | GACATAACTAATTACATGACTCGAGCTAATCAACATAGGTAAATATGGCC |
| HA-F520-F | TGCCGGATTATGCGTTCTCAATGAAAAACAACAAC |

| | |
|---|---|
| HA-F520-R | CATCATACGGATAATCTTGCATTGAATGTAAAGC |
| HA-STOP-F | TGCCGGATTATGCGTAGTCTGCTTTTTTACGTATATATATATATATATG |
| HA-STOP-R | CATCATACGGATAACTGACAATTAAATCGTTCAAC |
| HA-N671-F | TGCCGGATTATGCGAACAGCTGTTCAAAAAGC |
| HA-N671-R | CATCATACGGATAGCCATTTGTACCGCTATC |
| HA-K806-F | TGCCGGATTATGCGAAACAAGATAACAGAGACAAC |
| HA-K806-R | CATCATACGGATATTTGTCTATTTCCTTTGAATGAATG |
| NES_F | CTATGTTGATGGTGGCGGAAGCGGAGG |
| NES_R | GACATAACTAATTACATGACTCGAGCTAAATATCCAATCCGGCAAGTTTCAGG |
| NES | GGTGGCGGAAGCGGAGGGGGTTCAGGCGGTGGGAGTGGAGGTGGCTCCTTGGCTTTGAAGTTGGCTGGTCTTGACATCG GTTCTGGTGGTTCTGGTCTTGCCCTGAAACTTGCCGGATTGGATATTTAG |
| N527_F | AATAATTTGCCATCAATTAATCG |
| I335_R | GATCATAATAGGTGTTGTCG |
| Mga2-1113_F | TAGTCTGCTTTTTTACGTATATATATATATATATG |
| Mga2-626_R | ATCAACATAGGTAAATATGGC |
| Mga2-675_R | TGTGCTTTTTGAACAGC |
| Mga2-836_R | CGTACCGTTTTGGAATTC |
| Mga2-GFP_F1 | AGGGAACAAAAGCTGGAGCTCATCTTTGCTTCAAAGATTTTTTGC |
| Mga2_GFP_R1 | CTTTACTCATAACGAAATGTTCTGTTCG |
| Mga2_GFP_F2 | ACATTTCGTTATGAGTAAAGGAGAAGAAC |
| Mga2_GFP_R2 | CGAATTCCTGCAGCCCGGGGGATCCCTAACTGACAATTAAATCGTTC |
| GFP-1113_F | TAGGGATCCCCCGGG |
| GFP-626_R | ATCAACATAGGTAAATATGGCCTTTTTATTGTCATTGATC |
| Cy3-O2R_F | Cy3-TCGGACGTTGAACACTCAACAAACCTTATCTAGTGCCCAACCAGGTGTGC |
| Cy3-O2R_R | Cy3-GCACACCTGGTTGGGCACTAGATAAGGTTTGTTGAGTGTTCAACGTCCGA |
| O2R-Native_F | TCGGACGTTGAACACTCAACAAACCTTATCTAGTGCCCAACCAGGTGTGC |
| O2R-Native_R | GCACACCTGGTTGGGCACTAGATAAGGTTTGTTGAGTGTTCAACGTCCGA |
| O2R-Scrambled_F | TAGCATGCAACCGTCTCTAGTGTCACAACAGATACCCAGCGTAACTCAGT |
| O2R-Scrambled_R | ACTGAGTTACGCTGGGTATCTGTTGTGACACTAGAGACGGTTGCATGCTA |
| FAR_F | CCGGGGTTAGCGGGCCCAACAAAGGCGCTTATCTGGTGGGCTTCCGTAGA |
| FAR_R | TCTACGGAAGCCCACCAGATAAGCGCCTTTGTTGGGCCCGCTAACCCCGG |
| Ole1-qPCR_F | TGCTCTCTCTGGTAAAGTGCC |

| | |
|---|---|
| Ole1-qPCR_R | CACCACCGACAGCGTAGTAG |
| 5s_qPCR_F | GTTGCGGCCATATCTACCAGAAAG |
| 5s_qPCR_R | CGTATGGTCACCCACTACACTACT |
| URA3-qPCR_F | CATTGCGAAGAGCGACAAAGA |
| URA3-qPCR_R | ACCGGGTGTCATAATCAACCA |
| O2R-qPCR_F | AGTCGGCAGCTTTCTTTC |
| O2R-qPCR_R | CAAGACTCGTAGAAGCACAC |

**Table S2: plasmids used in the study**

| Number | Plasmid |
|---|---|
| 1 | pBEVY-GL HA-Rsp5 |
| 2 | pBEVY-GU 3xFlag-Mga2 |
| 3 | pET28-HIS-MBP-Mga2 (141-330) |
| 4 | pET28-HIS-MBP-Mga2 (141-330); K254R |
| 5 | pET28-HIS-MBP-Mga2 (141-330); R252A, R253A, K254A |
| 6 | pET28-MBP-3xFlag-Mga2(127-842); Δ371-529 |
| 7 | pET28-MBP-3xFlag-Mga2 (127-842); Δ371-529; K254R |
| 8 | pET28-MBP-3xFlag-Mga2(127-842); Δ371-529; R252A, R253A, K254A |
| 9 | pML104-2xHA Mga2 (ΔAnk) gRNA |
| 10 | pML104-2xHA Mga2 gRNA |
| 11 | pML104-2xHA Spt23 gRNA |
| 12 | pML104-Ole1 2xHA gRNA |
| 13 | pRS413-FAR (100bp)-URA3 |
| 14 | pRS413-O2R (50bp)-URA3 |
| 15 | pRS413-Ole1p (750bp)-URA3 |
| 16 | pRS416-2xHA Mga2-1-626 |
| 17 | pRS416-2xHA Mga2-1-626; Δ2-130 |
| 18 | pRS416-2xHA Mga2-1-626; L31E, L32E, L35E, L36E |
| 19 | pRS416-Flag Mga2 |
| 20 | pRS416-Flag Mga2; K980R, K983R, K985R |
| 21 | pRS416-Flag Mga2; L967A, P968A, Y870A |
| 22 | pRS416-Flag Mga2; R252A, R253A, K254A |
| 23 | pRS416-Flag Mga2; ΔAnk (626-836) |
| 24 | pRS416-TEF1-Flag Mga2-1-626 |
| 25 | pRS416-TEF1-Flag Mga2-1-626; K254R |
| 26 | pRS416-TEF1-Flag Mga2-1-626; R252A, R253A, K254A |
| 27 | pRS416-TEF1-Mga2 |
| 28 | pRS426-Flag-Mga2-(D519_HA_F520) |
| 29 | pRS426-Flag-Mga2-(S1113_HA_STOP) |
| 30 | pRS426-Flag-Mga2-HA (G670_HA_N671) |
| 31 | pRS426-Flag-Mga2-HA (K805_HA_K806) |
| 32 | pRS426-2xHA Mga2-1-626 |
| 33 | pRS426-2xHA Mga2-1-626; Δ2-130 |
| 34 | pRS426-2xHA Mga2-1-626; (LALKLAGLDI)X2 |
| 35 | pRS426-2xHA Mga2-1-626; L31E, L32E, L35E, L36E |
| 36 | pRS426-2xHA Mga2-1-626; K254R |
| 37 | pRS426-2xHA Mga2-1-626; K254R; Δ2-130 |
| 38 | pRS426-2xHA Mga2-1-626, K254R; Δ335-526 |

| | |
|---|---|
| 39 | pRS426-2xHA Mga2-1-626; K254R; (LALKLAGLDI)X2 |
| 40 | pRS426-2xHA Mga2-1-626; K254R; L31E, L32E, L35E, L36E |
| 41 | pRS426-2xHA Mga2-1-626; R252A, R253A, K254A |
| 42 | pRS426-Flag Mga2-1-626 |
| 43 | pRS426-Flag Mga2-1-626; K254R |
| 44 | pRS426-Flag Mga2-1-626; R252A, R253A, K254A |
| 45 | pRS426-Flag Mga2-1-675 |
| 46 | pRS426-GFP Mga2-1-626 |
| 47 | pRS426-GFP Mga2-1-626; K254R |
| 48 | pRS426-GFP Mga2-1-626; R252A, R253A, K254A |
| 49 | pRS426-GFP-Mga2-1-626; R252A, R253A, K254A; Δ2-130 |
| 50 | pRS426-GFP-Mga2-1-626; R252A, R253A, K254A; Δ335-526 |
| 51 | pRS426-GFP-Mga2 |
| 52 | pRS426-GFP-Mga2; R252A, R253A, K254A |
| 53 | Pk27-HIS-Sumo-Rsp5 |
| 54 | Pk27-HIS-Sumo-Ubc4 |

**Table S3: strains used in the study**

| Strain | genotype/description |
|---|---|
| BY4741 | MATa; his3-Δ1 leu2Δ0 met15-Δ0 ura3-Δ0 |
| *mga2* KO | BY4741; Δmga2:G418 |
| *spt23* KO | BY4741; Δspt23:G418 |
| 2xHA-Mga2 | BY4741 |
| 2xHA-Spt23 | BY4741 |
| 2xHA-Mga2 (ΔAnk) | BY4741 |
| *pdr5* KO | BY4741; Δpdr5:G418 |
| *mga2spt23* double deletion | MATα; his3-Δ1 leu2Δ0 lys2Δ0 ura3-Δ0; Δspt23:kanMX4: Δmga2:natMX |
| 4KO-LD; *mga2* KO | MATα; his3-Δ1 leu2Δ0 lys2Δ0 ura3-Δ0; are1::KanMX are2::KanMX lro1::loxP dga1::loxP; mga2Δ:hgh |
| NAB2-mCherry | MATa; his3-Δ1 leu2Δ0 met15-Δ0 ura3-Δ0; trp1Δ:natMX; NAB2-mCherry:TRP1 |
